# Eco-evolutionary feedback can stabilize diverse predator-prey communities

**DOI:** 10.1101/2022.07.29.502084

**Authors:** Stephen Martis

## Abstract

Ecological models with random interactions have provided insight into the problem of diversity, particularly showing that high variance in the distribution of interaction rates can lead to instability, chaos and extinction. However, these models have traditionally neglected evolution, which is central to the generation of biological variation and can act on timescales comparable to ecological change. We demonstrate that when a stochastic predator-prey system is coupled to high-dimensional evolutionary dynamics, high variance interactions counter-intuitively stabilize the population, delaying extinction and increasing the total population size. Using both stochastic and deterministic simulations and theory based on the statistical physics of disordered systems, this stabilizing effect is shown to be driven by an eco-evolutionary feedback loop which causes the population size to grow as a power law of the variance of the interactions. We show that the stable regime corresponds with the clonal interference regime of population genetics. We conjecture that qualitative aspects of our results generalize to other evolving complex systems.

## Introduction

The relationship between stability and diversity is a core problem in theoretical ecology^1^ – how can so much natural diversity (down to the lowest taxonomic levels^2^) persist in the face of strong competition, stochasticity, and potential chaos? And how does this phenotypic diversity contribute to the stability of natural populations? In order to address these questions, theorists have taken to analyzing model ecosystems with randomly drawn interactions^3–5^, which respect species heterogeneity without assuming any fine-tuned community structure. A key result from this long line of work is that ecosystems with interaction rates drawn from a broader distribution (i.e. ecosystems that are more diverse) are less likely to be stable^3,4^. However, these works have typically neglected the evolutionary process, which generates phenotypic diversity in the first place. Since evolution is coupled to the birth of new individuals, it can act on timescales comparable to ecological change, especially in larger natural populations. In this work, we show that when evolution acts sufficiently quickly, ecological and evolutionary processes can enter a feedback loop that stabilizes otherwise unstable population dynamics. Moreover, we show that in this eco-evolutionary context, sufficient phenotypic diversity is *required* for stability, in contrast to classical results.

Long ago, it was recognized that many-species Lotka-Volterra systems (with or without randomly drawn coefficients) readily admit a wide range of possible dynamics, including chaos^3,6^. Such chaotic dynamics can in turn lead to rapid cascades of extinctions^7^, that can be relevant at the ecosystem level. A key result, originating with May^3^ and further explored by others^4,5^, is that, for a broad class of disordered ecological models, the variance of the distribution of interaction rates controls the stability of the dynamics – the more varied the interactions, the less likely a large ecosystem will be stable. This suggests that phenotypic diversity negatively impacts ecological stability, as was recently demonstrated in a simplified laboratory setting^7^.

Over the years, there have been many attempts to propose mechanisms that might stabilize ecosystems with high variance interactions. These mechanisms, while plausible, are typically finely tuned, or have particular structural requirements, and as such are unlikely to generalize. Among those proposed are the ‘Kill-the-Winner’ hypothesis^8^, spatio-temporally fluctuating environmental conditions^9^, and higher-than-pairwise interactions^10^. Most recently, a storage effect-like mechanism generated by stable spatio-temporal chaos was proposed to promote diversity^11,12^. It was further shown that this type of spatiotemporal chaos can admit further diversification when evolution is vanishingly slow and when there is sufficient starting diversity^13^. However, this leaves open the question – how does such substantial initial diversity come to be in the first place^14^? Intriguingly, there has been some work to suggest that evolution might itself stabilize small or relatively simple ecological models^15–19^, while other work suggests that evolution can be destabilizing^20,21^, while still more work suggests that evolution might be either stabilizing or destabilizing depending on the context^22^. However, to the best of our knowledge, no work has connected evolution to the ‘large, complex’ regime when ecology and evolution act on comparable, shorter timescales.

In order to address this gap, we developed a fully stochastic eco-evolutionary predator-prey model with randomly drawn interactions. The underlying ecological interactions are defined in such a way that they facilitate unstable population dynamics over their entire parameter space – in a qualitative sense, they are ‘maximally’ unstable, resulting in rapid extinction of one or both species. They are also biologically-inspired, defined through a sequence-matching interaction, rather than statistically defined^11,12,23^. We couple these ecological dynamics to an evolutionary process on a high-dimensional genotype space. Using efficient stochastic simulations, deterministic simulations and theory based on the statistical physics of disordered systems, we show that in this context, evolution is able to stabilize the predator and prey population dynamics, but only when the variance of the interaction rates is sufficiently *high*, thus inverting the classical intuition. We show that in a population of discrete individuals, complex strain dynamics undergird this population-level stability. We also demonstrate that the transition to stability coincides with the clonal interference regime of population genetics. Finally, we discuss how our qualitative results might generalize to other eco-evolutionary scenarios. For the interested reader, the manuscript is organized such that the main results are presented in a more accessible manner in the main text, while technical details are contained in the Appendices.

### Model description

We consider a system of many predators and prey in which every predator-prey pair has a randomly drawn interaction rate. We assume nutrient saturation and well-mixed conditions with no explicit population size control for the predator or prey population. Our qualitative results will not appreciably change with the addition of carrying capacity (see Appendix F for a more explicit discussion). We also assume no spatial structure, which may also contribute to stabilizing the population dynamics^11,23^. All of these and more auxiliary processes might play some role in the long time behavior of real populations, but we make the choice to neglect them so that we can isolate the destabilizing role of diverse interactions. We discuss what sets the relative importance of these auxiliary processes and possible extensions of the model in Appendix F.

We also assume that the predation process is such that predators do not always increase their population size when they consume prey. We make this choice so that the interaction matrix is not anti-symmetric. Anti-symmetric many-strain predator-prey models, which possess a conserved quantity, are stably chaotic at the mean field level^24^, while many-strain predator-prey models with interactions that have broken anti-symmetry exhibit diverging chaos^11^. Such broken anti-symmetric interactions can be generated by biological phenomena like phage burst size or the generation of non-viable particles. Both of these processes create a discordance between the number of predation events and the number of viable predators that are born. We do not attempt to explicitly model the phenomena of burst size and non-viable particles, opting instead to focus on more generic aspects of predator-prey interactions without strict anti-symmetry.

The anti-symmetric component of the interaction between prey *i* (with abundance *X_i_*) and predator *k* (with abundance *Y_k_*) occurs with rate *a_ik_*. The *a_ik_* are strictly positive and drawn from a distribution with mean *a* and standard deviation *σ_a_*. In the anti-symmetric reaction a prey is consumed and an additional predator is born. The anti-symmetry breaking component occurs with strictly positive rate *s_ik_*, which is drawn from a distribution with mean *s* and standard deviation *σ_s_*. In the anti-symmetry breaking reaction a prey is consumed and no additional predators are born. A schematic of these interactions is illustrated in Fig. 1. Such ‘split’ interactions can be interpreted as a simplified model of variable burst size as discussed in Appendix A. In the simulations in the main text, for ease of presentation we set *σ_a_* = *σ_s_* = *σ* and *a* = *s* = 10^−3^. In the most general case, we assume that each prey has a birth rate, *b_i_* and each predator has a death rate, *d_k_*. In our simulations we set *b_i_* = *d_k_* = 1. We discuss modeling choices and variations of the model in the Appendices A and F.

**Figure 1:**
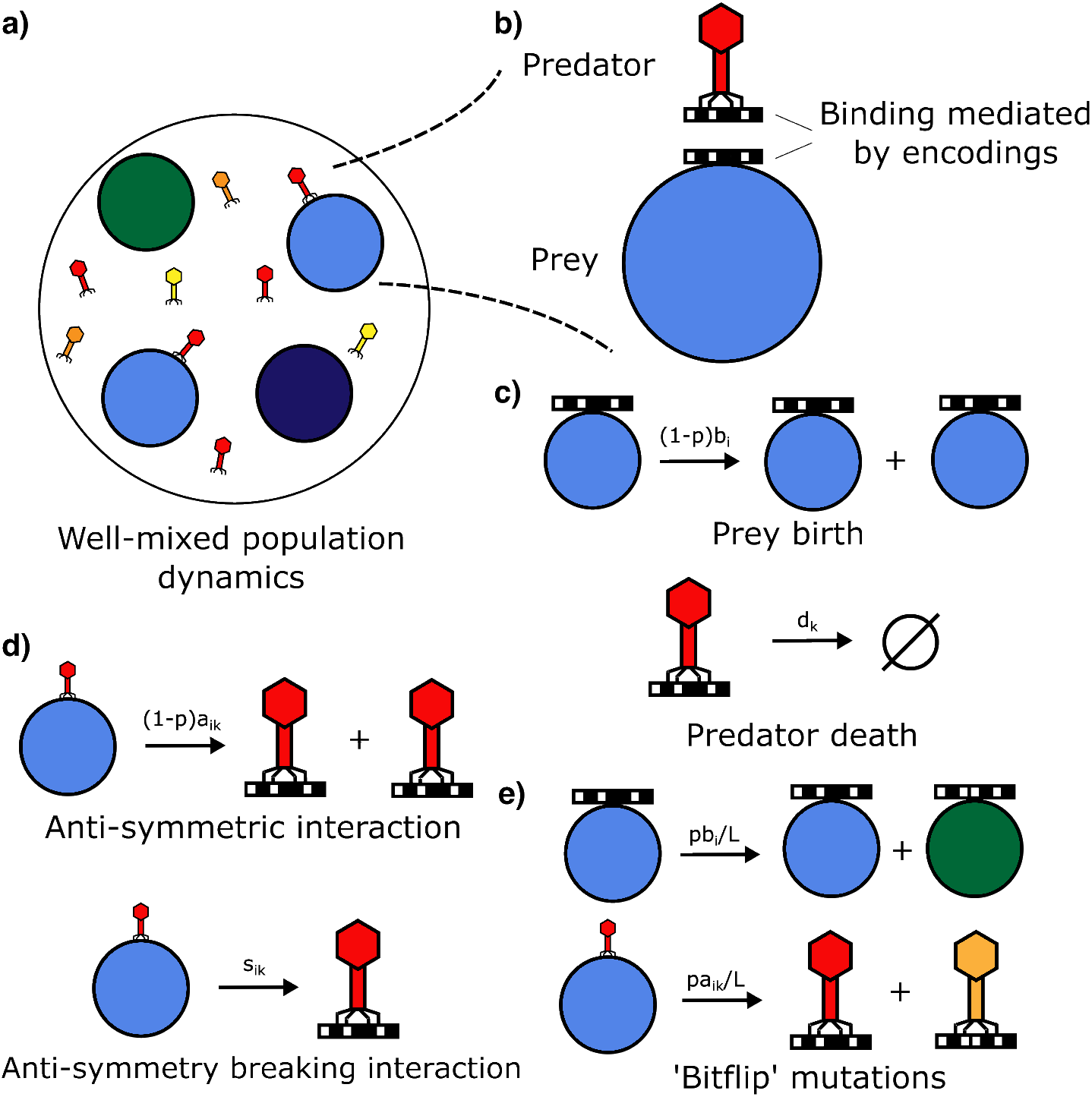
A schematic description of the population dynamic model we employ. a) The systems we consider are well-mixed and consist of two ‘species’ with many substrains. b) Interactions between the two types of species are mediated by bitstring ‘genotypes.’ Interaction rates are drawn from a distribution. c) Prey are born at a basal rate and predators die at a basal rate. d) Anti-symmetric interactions result in the death of a prey and the concurrent birth of an extra predator. Anti-symmetry breaking interactions result in the death of a prey and the maintenance of the predator population size. e) Mutations to the bitstring genotype probabilistically occur during events that result in the birth of a new individual. Note that there are key differences between the model and actual bacteria-phage systems, and the model should not be interpreted as one that will directly map onto a biological scenario. However, qualitative aspects of this model should generalize to more faithful representations of biological systems.

In addition, every predator and every prey is indexed by bitstrings of length *L_p_* and *L_b_*, respectively, so that there are 2*^L_p_^* predator and 2*^L_b_^* prey types in the model. In all that follows we consider *L_p_* = *L_b_* = *L*. We refer to the different values of the bitstrings as ‘genotypes.’ For each birth event (of predator and prey), there is a probability, *p*, that a point mutation occurs due to replication error, with probability 1 – *p* that a birth is clonal. We allow for mutations of ‘bitflip’ type so that only *L* mutant genotypes are accessible from a given genotype *i*. We denote this set of accessible genotypes 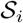. We present a schematic of these interactions in Fig 1. The mathematical form of the stochastic reactions is presented in a table in the Appendix A.2 and we describe the corresponding mean-field equations in the Methods.

The form of the interactions and the genotypes can be motivated biologically. Often interactions between cell types are mediated by binding between proteins. For instance, we can imagine a phage-bacteria system in which the phage must recognize and bind to a receptor on the surface of a bacterium in order to continue its life cycle. Or we might consider a T cell recognizing antigen. The bitstrings then represent an encoding of different sequence-specific receptors and ligands. The reaction rates vary because different pairs of binding site sequences have different binding affinities. Importantly, these rates are assumed fixed by the physics of the interactions between different receptor-ligand sequence pairs. We assume that the different genotypes are closely related enough that these binding affinities are drawn from a single distribution with a well-defined mean and variance. However, we allow that single mutations can vastly change the binding affinity of pairs of proteins, which has been observed in mutagenesis experiments^25,26^. In the main text, the interactions are drawn from a log-normal distribution, which can be motivated by Arrhenius type binding kinetics (Materials and Methods, Stochastic simulations). We discuss phylogenetically related interaction phenotypes in the Appendix F. However, they do not affect our qualitative observations.

### Population stability with large σ, large *p*

We can simulate the stochastic dynamics outlined above and plot the total prey and predator population sizes, *X_tot_* and *Y_tot_*, respectively. Populations are initialized to be monoclonal. Representative simulation trajectories of the total population sizes for *σ* = 1, *a* = 0.001 and *σ* = 0, *a* = 0.001 (*p* = 0.1 fixed) are shown in Fig. 2. In the neutral case, *σ* = 0, either the predator or prey population goes extinct on simulation relevant timescales due to demographic noise, which has been demonstrated before^16,27,28^. When we increase *σ*, we find the surprising result that population-level fluctuations are reduced, and extinction is less likely. We find that *σ* several orders of magnitude greater than the mean interaction strength damps population oscillations in both stochastic and deterministic mean-field simulations. Moreover, we find the counter-intuitive result that the total population sizes grow with increased disorder when compared to their neutral unstable fixed point values (Fig. 3b).

**Figure 2:**
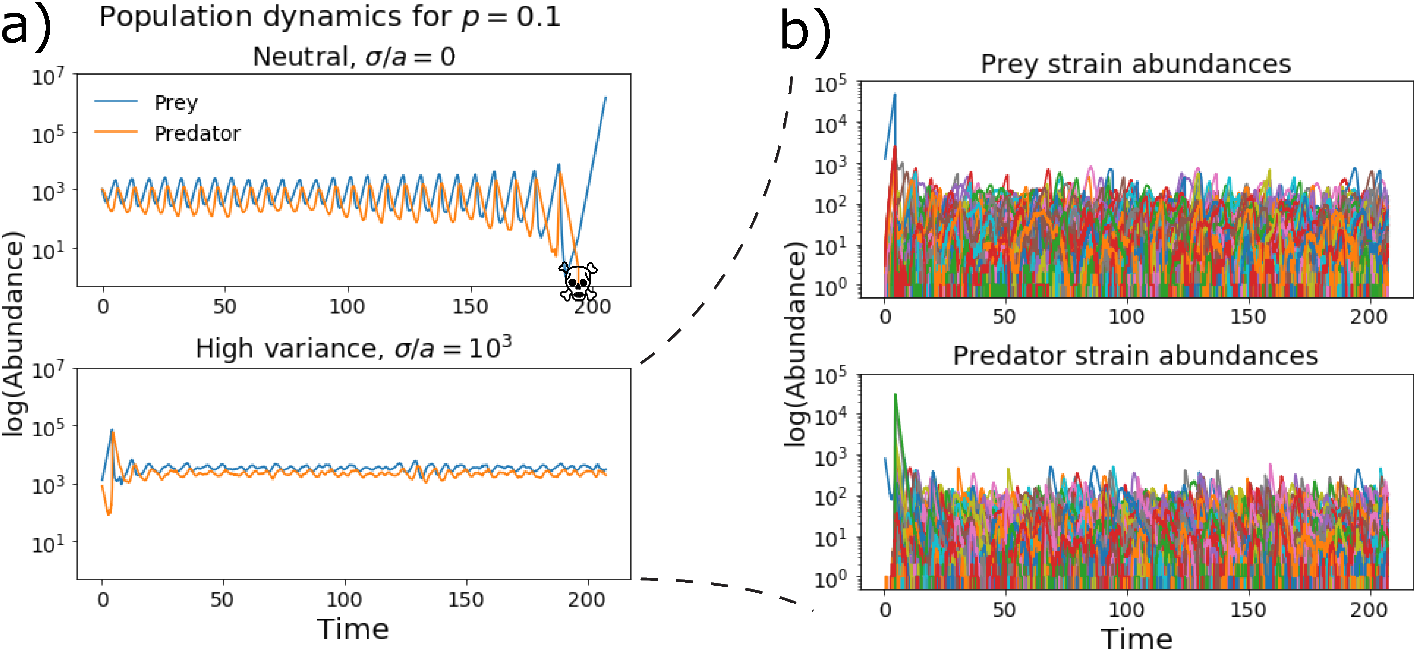
a) Stochastic simulations of the total predator and prey population size for different interaction spread, *σ* (plotted on a logarithmic scale). The neutral eco-evolutionary process (*σ_a_* = *σ_s_* = 0, *a* = *s* = 10^−3^) leads to rapid extinction of either the predator or both populations. Extinction always occurred in the neutral case for 100 runs with 2 million interactions. For highly variable interactions (*σ_a_* = *σ_s_* = 1, *a* = *s* = 10^−3^), we see that the dynamics are stabilized. Both simulations have mutation probability *p* = 0.1. Extinction never occurred in the high variance case in over 100 runs with 2 million interactions. b) The strain dynamics underlying the stable population dynamics can appear chaotic with constant extinction and recolonization by mutation.

**Figure 3:**
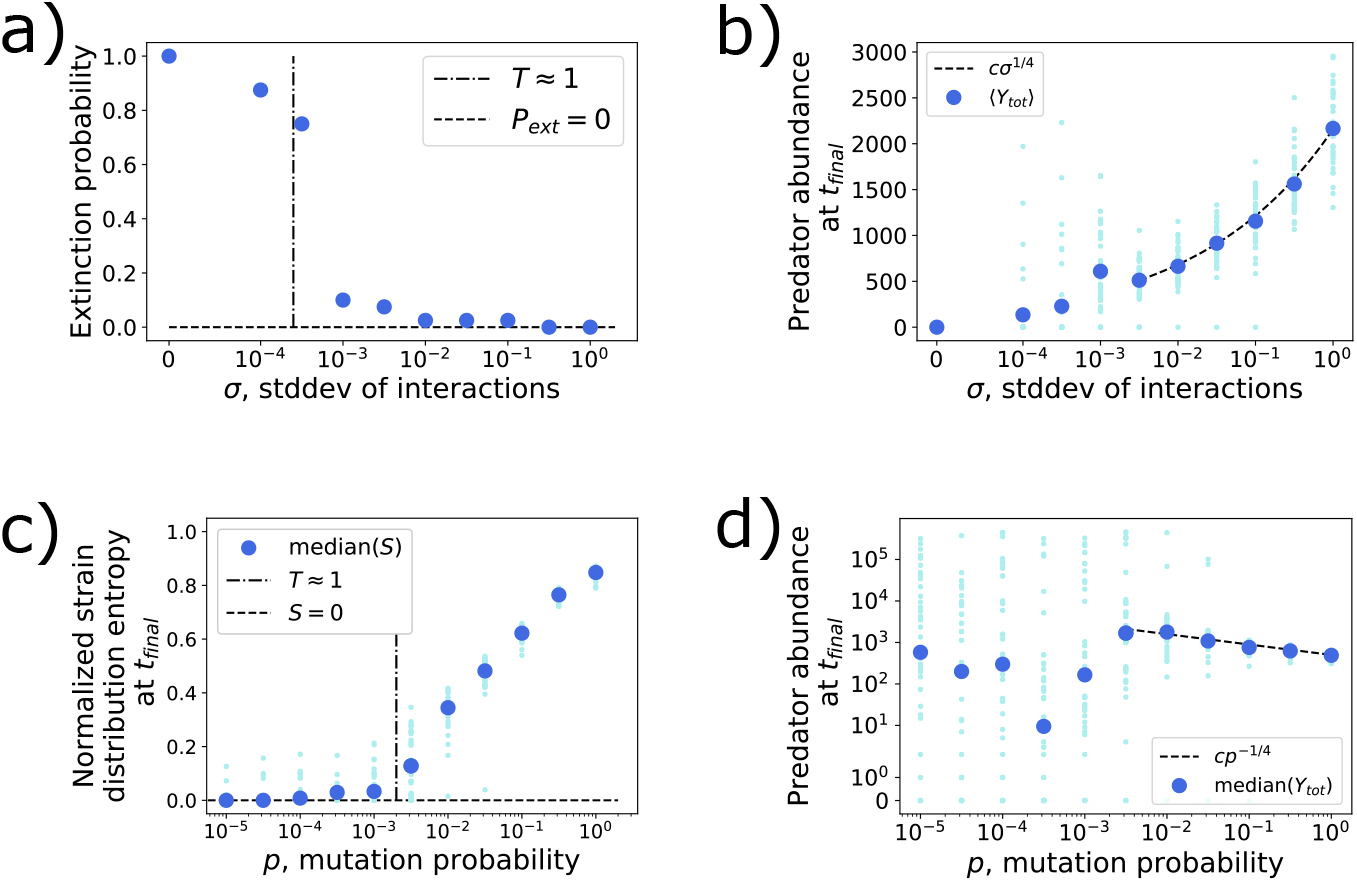
The transition between extinction and stability and the accompanying increase in population size. a) The extinction probability (blue circles) sharply drops when the parameter *T* (Eq. 2) is close to one. We expect many of the simulations to the left of the transition will go extinct given sufficiently long simulation time. b) The predator population size at the end of simulation time grows with the standard deviation of the interactions. Initialized with a predator-prey pair whose interactions are randomly drawn, log *a*_00_ ~ Normal(*a*, *σ_a_*), log *s*_00_ ~ Normal(*s*, *σ_s_*). Large dark blue dots are averages over simulations. Small light blue dots are individual simulations. All averages are over 50 simulations. Dashed line is a power-law fit. For both a) and b) *p* = 0.1. Though it is not shown, the prey population size grows with a similar exponent when conditioned on predator survival (if the predator goes extinct the prey population grows indefinitely). c) Entropy of the predator strain distribution (normalized by log(2*^L^*)) at the final time point of the simulation, varying the mutation probability, *p*. d) Predator abundance at the final time point of the simulation, varying the mutation probability, *p*. Large dark blue dots are averages over simulations. Small light blue dots are individual simulations. All averages are over 50 simulations. Dashed line is a power-law fit. The power law exponent has a similar magnitude to panel b). Initialized with a pair of strains whose interactions are randomly drawn, log *a*_00_ ~ Normal(*a*, 0.1), log *s*_00_ ~ Normal(*s*, 0.1). Because of the presence of long-lived, potentially metastable states (due to random initialization) we look at the median over realizations instead of the mean.

We also vary the mutation probability *p*, while holding *σ* = 0.1 fixed (Fig. 3c,d). We see, unsurprisingly, for very low mutation rates, that either predator or prey extinction is highly likely. However, at intermediate p, we see that sometimes the system stays in a low diversity state (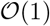 strains at a time) for long times. This happens by chance and depends on the random initial conditions and realization of the interactions. Such a population will still go extinct albeit on an anomalously long timescale (e.g. these cases appear to be metastable). As we increase *p*, amplitude fluctuations stabilize until we reach *p* = 1, when the population level dynamics fluctuate around a fixed point. While the *p* = 1 limit may not be natural (where all births result in mutants), it is a useful one to keep in mind when trying to understand the mechanism of the stabilizing feedback.

When examining the strain level dynamics, we see that strong fluctuations can undergird the population-level stability (Fig. 2b). Despite these complicated dynamics, the strain abundances remain finite and approximately uniform when averaged over very long timescales. The population-level stabilization effect is brought about by the interspecific feedback loop between the predator and prey strains. However, through the lens of dynamical mean-field theory (DMFT)^5,11^, this interspecific feedback can be cast as effective *intraspecific* interactions. In other contexts, explicit intraspecific interactions have been identified as a stabilizing mechanism^29,30^. Because evolution is also a relevant process in our model, intraspecific diversity is continually renewed so that stability can persist even in the face of individual strains going extinct. This process differs from bet-hedging or so-called ‘storage effect’ phenomena in the mechanism by which diversity persists. In these more classical modes of ‘ecological insurance,’ extinction is avoided through the presence of seed banks or spatial refugia. These refugia may even temporally fluctuate, as is the case in a recently examined class of spatially structured ecological models^11,23^. Whereas, in the present model, the insurance is dynamically and locally generated through the stochastic birth of mutants in a high-dimensional genotype space without the need to impose a spatial structure. Furthermore, in the eco-evolutionary context there is a balance whereby the intraspecific diversity is not so excessive that chaotic fluctuations drive population extinction (see Appendix D.1). We go on to discuss the details of the DMFT approach in the next section and in Appendix C.

### Total population sizes are stable in the many strain limit

From simulations, it is apparent that for sufficiently large *σ* and *p*, the total prey and predator population sizes, denoted *X_tot_* and *Y_tot_*, respectively, are approximately constant in time. Moreover, the averages of these abundances increase with increasing disorder, *σ*. In order to understand this behavior, it is instructive to look to the mean-field equations that describe the dynamics of *X_tot_* and *Y_tot_*. The total population size with evolution is governed by the same form of mean field equations as the total population size without evolution (i.e. summing over all types, the mutation terms cancel, see Methods).

We can then appeal to dynamical mean field theory (DMFT)^11,12,31^ to de-couple the interactions between predator and prey strains and replace them with effective ‘self-interactions’ when the genotype space is large. DMFT and similar methods have been successfully used to analyze highly diverse ecological scenarios without coupled evolutionary dynamics^4,5,11,12^. We find that in the limits (taken in order from first to last) *L* → ∞, *p* → 1, *t* → ∞, the mean field dynamics of the total predator and prey population sizes are quite simple (see Methods and Appendix C.2), achieving approximate homogeneity in strain abundances at long times. The disordered interactions generate feedback in the form of effective time-integrated response functions (referred to as static ‘susceptibilities’), denoted *χX* and *χY*. While obtaining closed form expressions of these quantities is difficult, they are readily measured with simulations. The prey susceptibility provides negative feedback on the predator population, and acts like a carrying capacity – when a predator is at high abundance, it depletes its preferred prey decreasing its growth rate in the future. Without mutation, the predator susceptibility should have the same sign. However, when the mutation rate is sufficiently high, the predator susceptibility provides positive feedback on the prey population – when a focal prey is at high abundance, it is consumed by its most avid predators which results in the birth of potentially less avid mutant predator offspring. When observed over the characteristic death rate of the predator, the focal prey’s growth rate net increases. This change in sign of the susceptibility due to mutation allows for a valid nontrivial fixed point of the predator and prey population sizes.

For mean-dominated interactions, 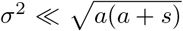, the susceptibilities are small and approximately constant so that the fixed point population sizes take on approximately constant values *X_tot_* ≈ 1/*a* and *Y_tot_* ≈1/(*a* + *s*) that would be expected from non-disordered interactions. For disorder-dominated interactions, 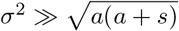, the product of the susceptibilities approaches 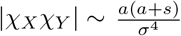. We make the ansatz that this approach is algebraic in the ratio *p*/*σ*. We confirm the leading term of our scaling ansatz by measuring the response in deterministic and stochastic simulations in the Appendix C.4. The divergence of the susceptibilities in turn causes the equilibrium population sizes to grow algebraically in the ratio *σ*/*p*, which is consistent with the measured population size over a broad range of *σ* in both models and a surprisingly broad range of *p* in the stochastic model (see Fig. 3):

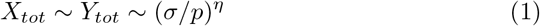

In the Appendix C.5, we argue that this power law exponent *η* is the generic result of the crossover from weak to strong disorder, rather than universal. The mean-field dynamics exhibit the power law growth of the population size in the strong disorder regimen only for *p* = 1 (Appendix B). However, the mean-field is unstable for *p* ≪ 1 without allowing for strain extinctions (see Appendix C.7). In the stochastic dynamics, such destabilizing effects of disorder are mitigated by demographic fluctuations and temporary extinction (Appendix D.1), while the susceptibilities, which are calculated from approximately uniform long-time strain abundances, exhibit analogous divergences with a different power law exponent. In this way, stochastic extinctions regularize divergent chaotic behavior that is present in the mean-field dynamics for *p* ≪ 1 while stabilizing *X_tot_* and *Y_tot_* over a broad parameter range.

Moreover, using our previous analyses, we can estimate the typical time for the predator population to go extinct and the typical time for an individual predator to mutate. By comparing the two, we are able to estimate the transition from likely predator extinction (the ‘extinct phase’) to long-time stability (the ‘stable phase’) in the stochastic model. The ratio of the two timescales can be conveniently expressed in terms of measurable parameters:

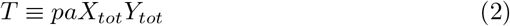

where *T* ≫ 1 is a necessary condition for stability. Using our scaling ansatz for the population sizes, we obtain consistent estimates for the transition point, up to 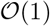 factors (see Figure 3b,d). We schematically map out the model’s phase diagram in log *p*-log *σ* space in Fig. 4. A more detailed discussion of the phase diagram, including a conjectured metastable phase, is contained in Appendix D.2.

**Figure 4:**
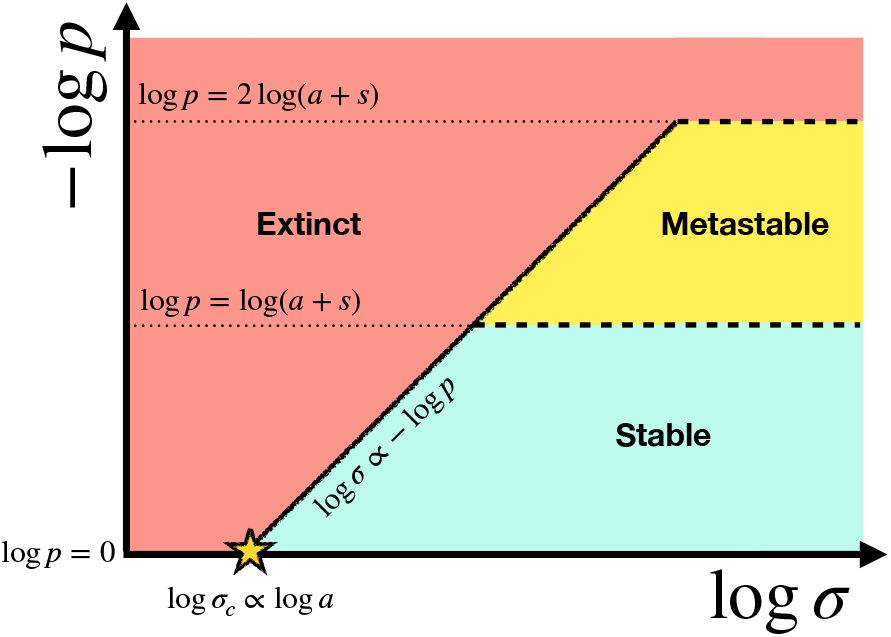
Proposed phase diagram of the extinct vs stable phases plotted in the log *p*-log *σ* plane for dynamics initialized with a single pair of strains and for sufficiently low *σ*. Note that the upwards direction corresponds to increasing – log *p* (which is the more natural scale since *p* ≤ 1). There is a point *σ* = *σ_c_* where the critical line intersects with the boundary, *p* = 1, which can be interpreted as the minimal diversity required to mitigate stochastic extinction. This point is predicted to scale with the mean interaction rate, a. For sufficiently high *σ* and log *p* ≥ log(*a* + *s*) the predator and prey populations are diverse and long-lived, i.e. ‘stable.’ The boundary of the stable region will expand (shift up and to the left) for larger populations (whose sizes are set by 1/*a* and 1/(*a* + *s*)). There is also a conjectured region where the long-time eco-evolutionary behavior is conjectured to be metastable, where the populations are potentially long-lived but not diverse. In this scenario one of the populations can go extinct on a longer timescale than the timescale of extinction in the extinct phase, but a shorter timescale than the timescale of extinction in the stable phase. This is discussed at length in the Appendix D.2. Panels a, c in Fig. 3 represent slices through this schematic phase diagram.

The quantity *T* can also be interpreted as the typical number of mutations produced by the predator population over a single generation (with the generation time set by the birth and death rates *b_i_* = *d_k_* = 1). Therefore stability requires:

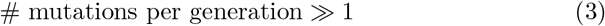

This condition corresponds with the well-known clonal interference regime from population genetics^32^, in which multiple mutants of differing fitnesses are simultaneously active within a population of fixed size. In turn, our results can be translated into the language of population genetics: clonal interference is crucial for the long term stability of large, well-mixed predator-prey populations.

However, we find that in the stable phase, appropriately defined strain fitness distributions do not have clear limiting shapes, which is inconsistent with traveling wave models that are associated with clonal interference and concurrent mutations in the literature^32–34^. This is despite the fact that similar eco-evolutionary models can be projected onto effective traveling wave models in suitable limits^17,35,36^ (see Appendix E for a more extended discussion). In the present model, the predator (prey) strain fitnesses are dependent on the composition of the prey (predator) population, and as such are dynamic quantities that can vary over orders of magnitude on timescales much shorter than extinction or mutation timescales, so the population fitness distribution is never able to ‘settle’ into a fixed shaped. Identifying whether a suitable limit exists whereby the present model might be projected onto an effective traveling wave model presents an interesting avenue for future work.

## Discussion

We have constructed a model of predator-prey co-evolution on a high dimensional genotype space. Using the model, we have demonstrated that, contrary to common intuition, highly disordered interactions serve to stabilize populations when mutations are sufficiently frequent. While the population level dynamics are stationary and resistant to stochastic extinction at long times, the strain level abundances can strongly fluctuate on short timescales when there is demographic noise. In the Appendix D.3, we discuss how these dynamics qualitatively resemble the so-called Griffiths phase phenomenon from statistical physics^37–39^, whereby the dynamics proceed through disjoint ‘rare’ regions of genotype space becoming active for finite characteristic times. We have proposed an eco-evolutionary ‘phase diagram’ and have provided a measurable parameter combination that can serve as an indicator of eco-evolutionary resilience in predator-prey populations. In addition, we have shown that stability corresponds with the well-characterized clonal interference regime from population genetics. However, population fitness distributions can be qualitatively different from established “traveling wave” theories^33^ of rapid adaptation when the underlying landscape of interaction rates is rugged.

We expect that our qualitative results should hold for other mutational architectures and for phylogenetically correlated phenotypes, so long as viable strains are sufficiently well-connected by mutation (see Appendix F). Furthermore, we expect that our qualitative results should hold in certain parameter regimes of other stochastic ecological models including ones that more closely mimic bacteria-phage interactions, that incorporate burst sizes, spatial structure, and other types of interactions. Given the generality of the stabilization mechanism, we also expect the stability results to be relevant in parameter regimes of other model systems where co-evolution is ubiquitous, including intrahost immune-pathogen dynamics^40^ and epidemiological models of individuals’ immune response^17^. Specifically, our theory is valid in certain asymptotic limits of our model. Therefore, we expect that our results should apply in the asymptotic limit of sufficiently similar models^41,42^, and so can be expected to be fairly general.

However, there remain many potential refinements and extensions. This is primarily a model of microdiversity, but diversity exists at many scales. In some cases, on longer timescales, innovations emerge to disrupt the ecological setting^43,44^, corresponding to changing the signs of the mean interactions in our model. Sometimes the appropriate scale is one at which localization^45^ or traveling waves^17,33^ might be the relevant phenomenology in phenotype space, rather than the phenomenology presently described. It would be exciting to directly connect the current framework to these and other important eco-evolutionary scenarios.

## Materials and Methods

### Mean-field equations

These stochastic dynamics lead to the following mean-field equations:

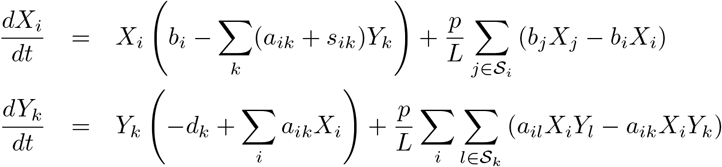

The first terms on the right hand side of each equation are identical to the terms of a standard mean-field predator-prey model, while the subsequent terms are novel and due to mutation. These mutation terms take the form of diffusion operators on a hypercube (the graph connecting the bitstring genotypes).

Above, we have assumed that the mutation probability, p, is constant across both predator and prey genotypes and that potential mutants are uniformly accessible, although both of these assumptions may be relaxed. Importantly, we assume that point mutation due to replication error is *explicitly coupled* to births, which in the case of the predator types only come about due to ecological interactions. A totally independent mutation rate, *μ*, can only be consistent when *p* is very small so that evolutionary processes are slow and act on statistically steady ecological timescales or if mutational processes unrelated to replication dominate (e.g. mutagens or horizontal gene transfer). We refer to such a model as the ‘*μ*-model.’ The *μ*-model, with 1/*μ* much greater than any ecological relaxation timescale, is the realm of certain regimes of classical population genetics^46^ and so-called adaptive dynamics^47^. We discuss models in which mutation is not correlated with birth in the Appendix F.2 (as opposed to our ‘*p*-model,’ which is discussed in the main text). The contents of this paper primarily deal with steady regimes in which evolutionary timescales are at least comparable to or faster than the timescale of extinction due to ecological forces.

Summing over strains, we can then get the mean field equations for the total population size:

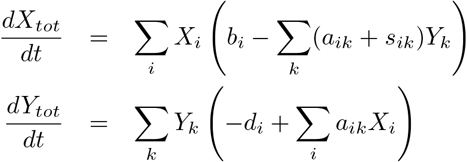

These are the equations we study in detail in the Appendix C and upon which we base our dynamical mean-field theory (DMFT) calculations. This analysis yields (in the *t* → ∞, *p* → 1, *L* → ∞ limit, applied from right to left):

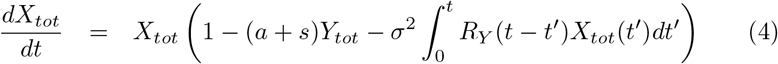

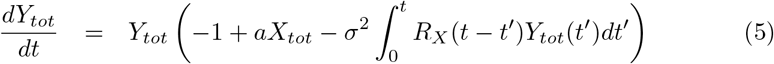

where *R_X_*(*t* – *t*′) and *R_Y_*(*t* – *t*′) are response functions that are defined self-consistently (see Appendix C).

### Stochastic simulations

We simulated the model using a version of Gibson and Bruck’s ‘Next Reaction Method’ (NRM)^48^, an exact stochastic simulation method with some performance advantages over the more commonly used Gillespie Algorithm (GA)^49^. The GA has an 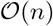 time complexity and while potentially faster in situations where the reaction propensities are multiscale (whereby one might order the vector of propensities in a clever way), our system does not exhibit clear multiscale behavior on long timescales. By exploiting data structures like priority queues and a reaction dependency graph, the NRM has a time complexity that scales as 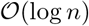 for a simulation with *n* reaction types. Therefore, it might provide substantial speedups when there are many reactions with sufficiently sparse dependencies, as is the case in our model. Since the NRM is an exact simulation scheme, there are no artefacts due to time discretization or other more ad hoc approximations (such as extinction cutoffs) and individual trajectories are guaranteed to be representative of the underlying stochastic process.

In our simulations, we set the prey birth rates and predator death rates to unity. We do not expect that slight departures from fixed birth and death rates will change our results significantly. We draw the interactions from a log-normal distribution with mean *μ* and standard deviation *σ*. In terms of traditional log-normal parameters, the mean and standard deviation of the exponentiated normal distribution, denoted *α* and *β,* respectively, we have:

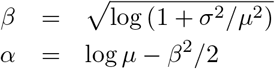

This choice guarantees that the total prey and predator populations will have 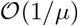 individuals in the mean field for sufficiently low *σ*.

We can motivate the choice of distribution physically, based on the picture of interactions mediated by binding due to protein contacts. In this picture, binding events will depend on the interaction energy of a pair of sequences of amino acids. For randomly drawn sequences, this gives an interaction energy which is the sum of many identically distributed site-specific energies, so that the binding energy distribution is Gaussian with some mean in the long sequence limit. The reaction rate can then be modeled as following an Arrhenius law, so that the distribution of rates follows a log-normal (with parameters discussed in the Methods). However, our qualitative results will be independent of our particular choice of distribution so long as it has a well-defined mean and variance.

We varied the width of the interaction rate distribution (*σ*) and the mutation probability (*p*) over many orders of magnitude. The maximal genome length we were able to achieve was *L* = 10, due to memory constraints. We ran simulations for 2 × 10^6^ reactions or until either the predator or prey class goes extinct, whichever came first. For simulations where strains are neutral with respect to each other (i.e. exchangeable, or *σ* = 0), 2 × 10^6^ reactions is sufficient to observe extinction with high probability.

### Deterministic simulations

We also simulated the deterministic mean-field dynamics. For these deterministic simulations, we implemented a 4th order Runge-Kutta scheme, with step size dependent on the number of strains, as determined by the genome length, *L*. For sufficiently many strains, the scheme was unstable unless prohibitively small step sizes were used. This can be understood from the fact that as more and more strains are added, the initial derivatives can become larger and larger so that smaller step sizes are needed to maintain numerical stability. Due to these issues, we were only able to simulate the deterministic model up to *L* = 10, with many plots being generated for *L* = 8 to save on compute. Comparisons to the stochastic model are discussed in the Appendices C and D.

## Code Availability

Relevant simulation code is available at https://github.com/stephen-martis/evolving-predator-prey.

## 1 Acknowledgments

I would like to thank members of the Hallatschek lab for comments on the many versions of this work, especially Oskar Hallatschek and Takashi Okada. In particular, I would like to thank Jonas Denk for in-depth comments. I would like to thank the Stanford Eco-Evo Journal Club, where discussions surrounding this work were initiated, and where an earlier draft of this work was critiqued. I would especially like to thank Benjamin H. Good, Daniel S. Fisher, Aditya Mahadevan, Gabriel Birzu and others for conversations about earlier drafts of this work. I would like to thank Benjamin Greenbaum for helpful feedback. This research used resources of the National Energy Research Scientific Computing Center (NERSC), a U.S. Department of Energy Office of Science User Facility located at Lawrence Berkeley National Laboratory, operated under Contract No. DE-AC02-05CH11231 using NERSC award BER-ERCAP0019907.

## References

1. McCann, K. S. The diversity–stability debate. Nature 405, 228–233 (2000).

2. Kashtan, N. et al. Single-cell genomics reveals hundreds of coexisting sub-populations in wild Prochlorococcus. Science 344, 416–420 (2014).

3. May, R. M. Will a large complex system be stable? Nature 238, 413–414 (1972).

4. Bunin, G. Ecological communities with Lotka-Volterra dynamics. Physical Review E 95, 042414 (2017).

5. Roy, F., Biroli, G., Bunin, G. & Cammarota, C. Numerical implementation of dynamical mean field theory for disordered systems: Application to the Lotka–Volterra model of ecosystems. Journal of Physics A: Mathematical and Theoretical 52, 484001 (2019).

6. Smale, S. On the differential equations of species in competition. Journal of Mathematical Biology 3, 5–7 (1976).

7. Hu, J., Amor, D. R., Barbier, M., Bunin, G. & Gore, J. Emergent phases of ecological diversity and dynamics mapped in microcosms. Science 378, 85–89 (2022).

8. Thingstad, T. F. Elements of a theory for the mechanisms controlling abundance, diversity, and biogeochemical role of lytic bacterial viruses in aquatic systems. Limnology and Oceanography 45, 1320–1328 (2000).

9. Hutchinson, G. E. The paradox of the plankton. The American Naturalist 95, 137–145 (1961).

10. Bairey, E., Kelsic, E. D. & Kishony, R. High-order species interactions shape ecosystem diversity. Nature communications 7, 1–7 (2016).

11. Pearce, M. T., Agarwala, A. & Fisher, D. S. Stabilization of extensive fine-scale diversity by ecologically driven spatiotemporal chaos. Proceedings of the National Academy of Sciences (2020).

12. Roy, F., Barbier, M., Biroli, G. & Bunin, G. Can endogenous fluctuations persist in high-diversity ecosystems? arXiv preprint arXiv:1908.03348 (2019).

13. Mahadevan, A., Pearce, M. T. & Fisher, D. S. Spatiotemporal Ecological Chaos Enables Gradual Evolutionary Diversification Without Niches or Tradeoffs. bioRxiv (2022).

14. Rubin, I., Ispolatov, Y. & Doebeli, M. Ecological diversity exceeds evolutionary diversity in model ecosystems. bioRxiv (2022).

15. Schenk, H., Schulenburg, H. & Traulsen, A. How long do Red Queen dynamics survive under genetic drift? A comparative analysis of evolutionary and eco-evolutionary models. BMC evolutionary biology 20, 1–14 (2020).

16. Xue, C. & Goldenfeld, N. Coevolution maintains diversity in the Stochastic “Kill the Winner” Model. Physical review letters 119, 268101 (2017).

17. Yan, L., Neher, R. A. & Shraiman, B. I. Phylodynamic theory of persistence, extinction and speciation of rapidly adapting pathogens. Elife 8, e44205 (2019).

18. Ackland, G. & Gallagher, I. Stabilization of large generalized Lotka-Volterra foodwebs by evolutionary feedback. Physical review letters 93, 158701 (2004).

19. Kondoh, M. Foraging adaptation and the relationship between food-web complexity and stability. Science 299, 1388–1391 (2003).

20. Shoresh, N., Hegreness, M. & Kishony, R. Evolution exacerbates the paradox of the plankton. Proceedings of the National Academy of Sciences 105, 12365–12369 (2008).

21. Cortez, M. H., Patel, S. & Schreiber, S. J. Destabilizing evolutionary and eco-evolutionary feedbacks drive empirical eco-evolutionary cycles. Proceedings of the Royal Society B 287, 20192298 (2020).

22. Abrams, P. A. The evolution of predator-prey interactions: theory and evidence. Annual Review of Ecology and Systematics, 79–105 (2000).

23. Roy, F., Barbier, M., Biroli, G. & Bunin, G. Complex interactions can create persistent fluctuations in high-diversity ecosystems. PLoS computational biology 16, e1007827 (2020).

24. Goel, N. S., Maitra, S. C. & Montroll, E. W. On the Volterra and other nonlinear models of interacting populations. Reviews of modern physics 43, 231 (1971).

25. Cunningham, B. C. & Wells, J. A. High-resolution epitope mapping of hGH-receptor interactions by alanine-scanning mutagenesis. Science 244, 1081–1085 (1989).

26. Lite, T.-L. V. et al. Uncovering the basis of protein-protein interaction specificity with a combinatorially complete library. Elife 9, e60924 (2020).

27. Parker, M. & Kamenev, A. Extinction in the Lotka-Volterra model. Physical Review E 80, 021129 (2009).

28. Parker, M. & Kamenev, A. Mean extinction time in predator-prey model. Journal of Statistical Physics 141, 201–216 (2010).

29. Yodzis, P. The stability of real ecosystems. Nature 289, 674–676 (1981).

30. Allesina, S., Miller, Z. R. & Servan, C. A. M. Intraspecific variation stabilizes classic predator-prey dynamics. bioRxiv (2021).

31. Mézard, M., Parisi, G. & Virasoro, M. A. Spin glass theory and beyond: An Introduction to the Replica Method and Its Applications (World Scientific Publishing Company, 1987).

32. Good, B. H., Rouzine, I. M., Balick, D. J., Hallatschek, O. & Desai, M. M. Distribution of fixed beneficial mutations and the rate of adaptation in asexual populations. Proceedings of the National Academy of Sciences 109, 4950–4955 (2012).

33. Desai, M. M. & Fisher, D. S. Beneficial mutation–selection balance and the effect of linkage on positive selection. Genetics 176, 1759–1798 (2007).

34. Fisher, D. S. Asexual evolution waves: fluctuations and universality. Journal of Statistical Mechanics: Theory and Experiment 2013, P01011 (2013).

35. Rouzine, I. M. & Rozhnova, G. Antigenic evolution of viruses in host populations. PLoS pathogens 14, e1007291 (2018).

36. Marchi, J., Lässig, M., Walczak, A. M. & Mora, T. Antigenic waves of virus–immune coevolution. Proceedings of the National Academy of Sciences 118, e2103398118 (2021).

37. Munoz, M. A., Juhász, R., Castellano, C. & Ódor, G. Griffiths phases on complex networks. Physical review letters 105, 128701 (2010).

38. Martin, P. V., Bonachela, J. A., Levin, S. A. & Muñoz, M. A. Eluding catastrophic shifts. Proceedings of the National Academy of Sciences 112, E1828–E1836 (2015).

39. Vojta, T. Rare region effects at classical, quantum and nonequilibrium phase transitions. Journal of Physics A: Mathematical and General 39, R143 (2006).

40. Nourmohammad, A., Otwinowski, J. & Plotkin, J. B. Host-pathogen co-evolution and the emergence of broadly neutralizing antibodies in chronic infections. PLoS genetics 12, e1006171 (2016).

41. Kadanoff, L. P. Scaling and universality in statistical physics. Physica A: Statistical Mechanics and its Applications 163, 1–14 (1990).

42. Hinrichsen, H. Non-equilibrium critical phenomena and phase transitions into absorbing states. Advances in physics 49, 815–958 (2000).

43. Good, B. H., Martis, S. & Hallatschek, O. Directional selection limits diversification and promotes ecological tinkering in the competition for many resources. PNAS (2018).

44. Blount, Z. D., Borland, C. Z. & Lenski, R. E. Historical contingency and the evolution of a key innovation in an experimental population of Escherichia coli. Proceedings of the National Academy of Sciences 105, 7899–7906 (2008).

45. Eigen, M., McCaskill, J. & Schuster, P. Molecular quasi-species. The Journal of Physical Chemistry 92, 6881–6891 (1988).

46. Gillespie, J. H. Some properties of finite populations experiencing strong selection and weak mutation. The American Naturalist 121, 691–708 (1983).

47. Geritz, S. A., Metz, J. A., Kisdi, É. & Meszéna, G. Dynamics of adaptation and evolutionary branching. Physical Review Letters 78, 2024 (1997).

48. Gibson, M. A. & Bruck, J. Efficient exact stochastic simulation of chemical systems with many species and many channels. The journal of physical chemistry A 104, 1876–1889 (2000).

49. Gillespie, D. T. Exact stochastic simulation of coupled chemical reactions. The journal of physical chemistry 81, 2340–2361 (1977).

