## Supplementary material for "Eco-evolutionary feedback can stabilize diverse predator-prey communities"

### Supplementary information for 'Eco-evolutionary feedback can stabilize diverse predator-prey communities'

Stephen Martis[1, 2]

[1]Current Affiliation: Computational Oncology, Department of Epidemiology & Biostatistics, Memorial Sloan Kettering Cancer Center, New York, NY, USA
[2]Physics Department, UC Berkeley, Berkeley, CA, USA

June 2022

### Appendix A Model details

#### A.1 Predator-prey vs. generalized Lotka-Volterra models

It is reasonable to model an ecological system as a generalized Lotka-Volterra model (GLVM), with complex interlocking foodwebs, but it is less clear how this type of landscape would fit in to an eco-evolutionary system of the sort that we present, which models fine-scale within-species diversity. Within a random GLVM architecture, by allowing for mutations between strains, it would be possible for a prey individual to give birth to its own predator. This type of process is unlikely on the relatively short timescales represented by our model. Studying a model in which the sign of a mutant's interactions can differ from its parent's is potentially interesting from a theoretical perspective, but care would need to be made to model this in a way that is biologically relevant. It would certainly be interesting to understand the eco-evolutionary process for more explicitly modeled interactions, which is conceptually possible within the current framework.

#### A.2 Stochastic reaction model

The reactions in our stochastic model are described in Table 1.

We can motivate the interactions in our model through the lens of receptor-ligand binding kinetics. This can be cast as a string matching problem whereby the 'right' pairs of strings have a low energy (and a high interaction rate) and the 'wrong' pairs of strings have a large binding energy (and low interaction

| Reaction | Description |
|---|---|
| $X_i \xrightarrow{(1-p)b_i} 2X_i$ | Clonal prey birth |
| $Y_k \xrightarrow{d_k} \emptyset$ | Predator death |
| $X_i + Y_k \xrightarrow{s_{ik}} Y_k$ | 'Lossy' predation (no birth) |
| $X_i + Y_k \xrightarrow{(1-p)a_{ik}} 2Y_k$ | Predation with clonal predator birth |
| $X_i \xrightarrow{pb_i/L} X_i + X_j \quad \forall j \in \mathcal{S}_i$ | Prey birth with random mutation |
| $X_i + Y_k \xrightarrow{pa_{ik}/L} Y_k + Y_l \quad \forall l \in \mathcal{S}_k$ | Predation with predator birth and random mutation |

Table 1: The set of reactions in our stochastic predator prey model.

rates). A model of the form we present would simulate an evolution experiment given binding measurements for pairs of predator and prey types.

The 'split' form of the interactions is primarily a mathematical convenience (into $a_{ik}$ and $s_{ik}$), but can also be interpreted as a weak form of variable burst size. In this interpretation the burst size is either two or one, with probabilities:

$$P(1) = \frac{a_{ik} + s_{ik}}{2a_{ik} + s_{ik}} \qquad P(2) = \frac{a_{ik}}{2a_{ik} + s_{ik}}$$

The mean burst size for the prey $i$-predator $k$ pair is then just $\frac{3a_{ik}+s_{ik}}{2a_{ik}+s_{ik}}$.

### Appendix B Qualitative agreement between stochastic and deterministic simulations

#### B.1 Population dynamics without mutation are seemingly chaotic with many effective extinctions

We simulate the population dynamics without mutation. This results in seemingly chaotic dynamics at the population level. At the strain level, there are wild fluctuations on a log scale (down to miniscule fractional abundances).

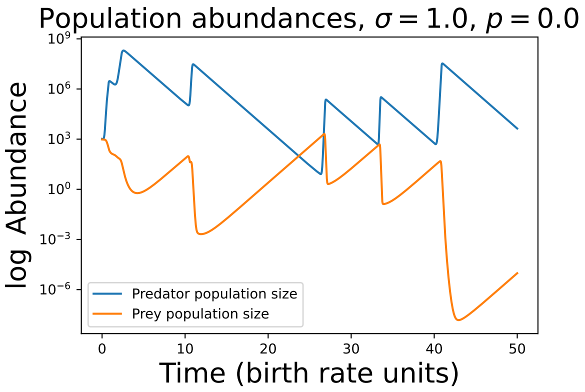


Figure 1: Prey and predator population size over time without evolution for $L = 8$. Notice the strong log-scale fluctuations.

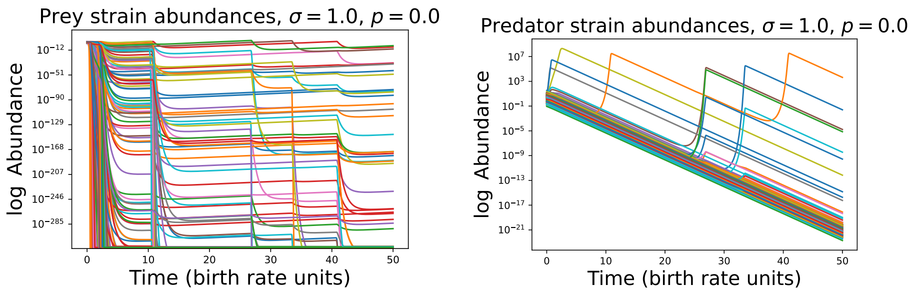


(a) (b)

Figure 2: (a) Prey and (b) predator strain abundances over time without evolution for $L = 8$. Notice the strong, unpredictable fluctuations, on a log scale, in addition to many apparent 'extinctions' (strain abundances decaying exponentially quickly).

#### B.2 Stability at the strain and population level with evolution

We simulated the deterministic dynamics with random coefficients without an extinction process. We fix $L = 8$, $p = 1$, $a = s = 10^{-3}$ and vary $\sigma$. We find that the deterministic simulations qualitatively agree with elements of the stochastic simulations. First we find apparent damped oscillations to a fixed point for large $\sigma$ when initialized near uniformity. As we increase $\sigma$, the population size of the predator and prey at the fixed point increases as a power of $\sigma$. We find that the population size grows approximately as a power law $X_{tot} \sim Y_{tot} \sim \sigma^{\eta}$, with $\eta \approx 1/2$. This power differs from the one we find in the stochastic simulations ($\eta = 1/4$). We expect that this is primarily due to demographic fluctuations, which induce a so-called 'Griffiths phase' in the stochastic simulations. We discuss this in a later section.

At the strain level, with mutation, the dynamics appear stable at long times (Figs. 4 and 5). For fixed $L = 8$ and weaker disorder (Fig. 4), the distribution of long time strain frequencies is tighter that for stronger disorder (Fig. 5). We discuss the distribution of the strain log frequencies in a future section.

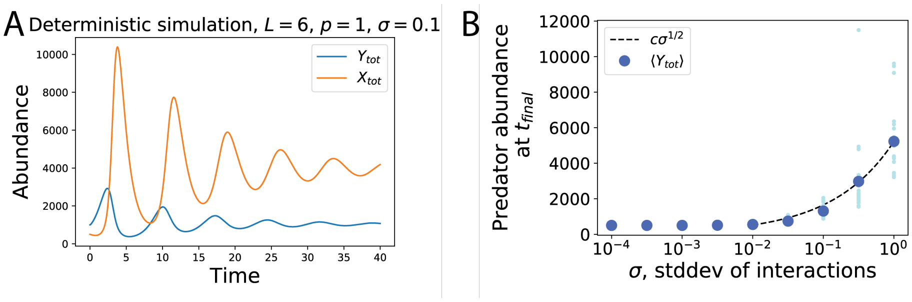


Figure 3: A.) Damped oscillations for a random deterministic simulation for $L = 6$, $p = 1$, and $\sigma = 0.1$. Notice that the total population size is larger than that which would be expected from the neutral case, $Y_{tot} = 500$, $X_{tot} = 1000$. The damped oscillations seem to persist for much lower $p$ and for much longer $t$, however the low $p$ simulations are too resource intensive to reach sufficiently long timescales while maintaining numerical accuracy. B.) For fixed $L = 8$ and $p = 1$, the final abundance increases with the variance of the interactions. The characteristic exponent is different from the stochastic case.

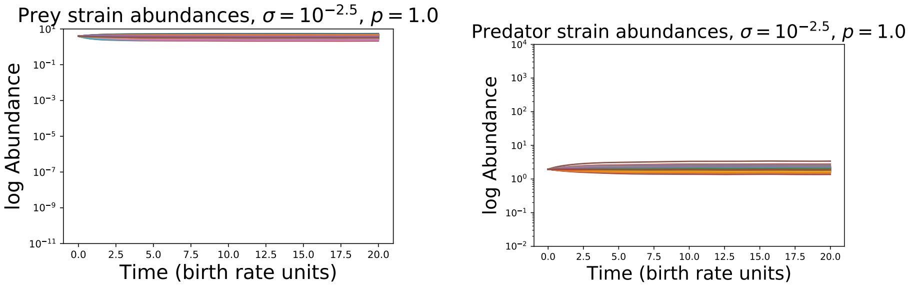


(a) (b)

Figure 4: (a) Prey and (b) predator strain abundances over time with evolution for $L = 8$ in the weak disorder regime ($\sigma = 10^{-2.5}$). Plotted on the same y-scale as the strong disorder plot in Fig. 5. Notice the apparent stabilization at long times, and the much tighter distribution of abundances. Any fluctuations at long times are even smaller than in the strong disorder case.

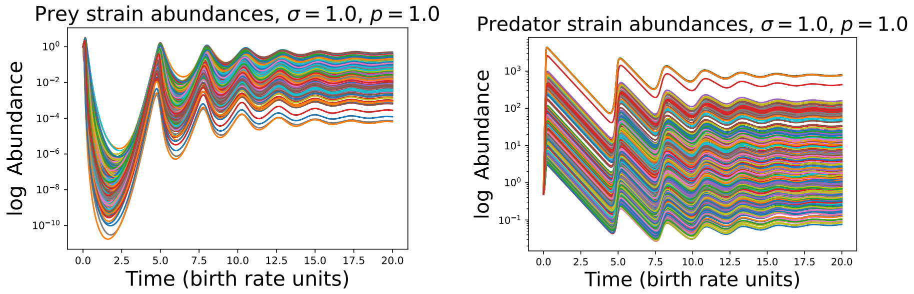


(a) (b)

Figure 5: (a) Prey and (b) predator strain abundances over time with evolution for $L = 8$ in the strong disorder regime ($\sigma = 1.0$). Notice the apparent stabilization at long times (despite some initial transients). Any fluctuations at long times appear to be bounded on a logarithmic scale, which is in stark contrast to the case without evolution. However, we see that for finite $L$, the distribution of long-time strain abundances is quite broad. Despite this, we provide evidence that, as $L \to \infty$, the long-time distribution of strains converges to a peak.

### Appendix C Dynamical mean field theory (DMFT)

#### C.1 DMFT equations for the total population sizes

Since there is agreement between stochastic and deterministic simulations, we work with the deterministic case to simplify our analysis and understand the mechanism of stabilization and the origin of power law growth in population size. We exploit the fact that the form of the dynamical equations for the total population size are identical regardless of whether or not mutation is present. We then use dynamical mean-field theory to get the equilibrium properties of the total population size.

We work in the case where $b_i = d_k = 1$. Since we are concerning ourselves with the mean-field limit, we have that there are no extinctions in this setting. Formally, we work in the large but finite $2^L$ limit, leaving in these factors for clarity about scaling and the relative size of different quantities. Later on, we take the thermodynamic limit, $L \to \infty$. We have that the mean field equations for the total abundances are given by:

$$\begin{aligned} \frac{dX_{tot}}{dt} &= \sum_i X_i \left( 1 - (a+s)Y_{tot} - \sum_{k=1}^{2^L} (\delta a_{ik} + \delta s_{ik}) Y_k + h_i(t) \right) \\ \frac{dY_{tot}}{dt} &= \sum_k Y_k \left( -1 + aX_{tot} + \sum_{i=1}^{2^L} \delta a_{ik} X_i + h_k(t) \right) \end{aligned}$$

where we have split the interactions into mean and fluctuating components:

$$\begin{aligned} a_{ik} &= a + \delta a_{ik} \\ s_{ik} &= s + \delta s_{ik} \end{aligned}$$

We have also added response fields $h_i$ and $h_k$ for convenience, which we will later set to 0. We can use the lens of dynamical mean-field theory to understand the properties of these quantities. The essence of dynamical mean field theory is that the sums over interactions:

$$\sum_k (\delta a_{ik} + \delta s_{ik}) Y_k \quad \text{and} \quad \sum_i \delta a_{ik} X_i$$

can be averaged over and treated self-consistently in the many-strain limit. In the language of statistical physics, the interactions are decoupled and replaced with an effective noise and 'self-interaction.' The dictionary for the mapping (which has been derived in other contexts [9, 32, 28, 19]) is that:

$$\begin{aligned} \sum_k (\delta a_{ik} + \delta s_{ik}) Y_k &\to (\sigma_a + \sigma_s) 2^{L/2} \eta_i^X(t) + \sigma_a^2 2^L \int_0^t R_Y(t,t') X_i(t') dt' \\ \sum_i \delta a_{ik} X_i &\to \sigma_a 2^{L/2} \eta_k^Y(t) - \sigma_a^2 2^L \int_0^t R_X(t,t') Y_k(t') dt' \end{aligned}$$

where we use the natural scaling suggested in [28] and carefully keep track of $2^L$ prefactors and minus signs. Note: there are technically $\mathcal{O}(2^{L/2})$ error terms, $\epsilon_i(t)$ and $\epsilon_k(t)$ in this mapping, that become irrelevant when we take the thermodynamic limit later on. For ease of presentation, we neglect them at this stage in the derivation, since they are subleading when compared to the noise and to the response.

In lieu of repeating the explicit derivations from [9, 32, 28, 19] and others, we provide an intuitive description of the two terms in each of these mappings. The first are zero-mean noises with correlation structure:

$$\begin{aligned}\overline{\eta_i^Y(t)\eta_i^Y(t')} &= 2^{-L}\sum_k Y_k(t)Y_k(t') \equiv C_Y(t,t') \\ \overline{\eta_k^X(t)\eta_k^X(t')} &= 2^{-L}\sum_i X_i(t)X_i(t') \equiv C_X(t,t')\end{aligned}$$

which is demanded by self-consistency. This term determines the invasion properties of the different strains (i.e. it contributes to strain $i$'s fitness while rare). When a strain is rare, all the strains it interacts with are approximately independent random variables, that are importantly approximately independent from the focal strain itself (since it is at low abundance).

The second term is the linear response when $X_i$ or $Y_k$ gets large. Intuitively this accounts for the fact that when $X_i$ large, it will perturb the $Y_k$ (which will result in births of $Y_k$), which will in turn feed back on $X_i$, causing its birth rate to decline in the absence of mutations. Similarly $Y_k$ at high abundance will deplete its preferred prey and reduce its birth rate in the future. The intuition is that, without mutation, feedback effects result in effective time-dependent carrying capacities for the predator and prey populations. Although, as we will see in the next section, mutation can change this intuition. The response functions are given (once again, due to self-consistency) by:

$$\begin{aligned}R_Y(t,t') &= \overline{\frac{\delta Y_k(t)}{\delta h_k(t')}}\Bigg|_{h_k=0} = 2^{-L}\sum_k \frac{\delta Y_k(t)}{\delta h_k(t')}\Bigg|_{h_k=0} \\ R_X(t,t') &= \overline{\frac{\delta X_i(t)}{\delta h_i(t')}}\Bigg|_{h_i=0} = 2^{-L}\sum_i \frac{\delta X_i(t)}{\delta h_i(t')}\Bigg|_{h_i=0}\end{aligned}$$

where these derivatives are in the functional sense.

Summing over all strains, assuming we are in a stable phase (where all strains might be seeded by mutation, so none go extinct indefinitely), we have the DMFT equations for the total population sizes:

$$\begin{aligned}\frac{dX_{tot}}{dt} &= \sum_i X_i\Big(1-(a+s)Y_{tot}-(\sigma_a+\sigma_s)2^{L/2}\eta_i^Y(t)\\ &\qquad -\sigma_a^2 2^L\int_0^t R_Y(t,t')X_i(t')dt'\Big)\\ \frac{dY_{tot}}{dt} &= \sum_k Y_k\Big(-1+aX_{tot}+\sigma_a 2^{L/2}\eta_k^X(t)\\ &\qquad -\sigma_a^2 2^L\int_0^t R_X(t,t')Y_k(t')dt'\Big)\end{aligned}$$

where we have set the response fields to zero now that they are no longer needed. We can rewrite these equations:

$$\begin{aligned}\frac{dX_{tot}}{dt} &= X_{tot}\Big(1-(a+s)Y_{tot}-(\sigma_a+\sigma_s)2^{L/2}\left\langle\eta^Y(t)\right\rangle_{X(t)}\\ &\qquad -\sigma_a^2 2^L\left\langle\int_0^t R_Y(t,t')X(t')dt'\right\rangle_{X(t)}\Big)\\ \frac{dY_{tot}}{dt} &= Y_{tot}\Big(-1+aX_{tot}+\sigma_a 2^{L/2}\left\langle\eta^X(t)\right\rangle_{Y(t)}\\ &\qquad -\sigma_a^2 2^L\left\langle\int_0^t R_X(t,t')Y(t')dt'\right\rangle_{Y(t)}\Big)\end{aligned}$$

The angle brackets denote the (grand canonical) average:

$$\langle\mathcal{O}(t)\rangle_{Y(t)}=\frac{1}{Y_{tot}(t)}\sum_k\mathcal{O}_k(t)Y_k(t)$$

which is formally different from the overline (microcanonical) average:

$$\overline{\mathcal{O}}=2^{-L}\sum_k\mathcal{O}_k$$

We finally assume we are in a statistically steady state so that correlation and response are only a function of time differences:

$$\overline{\eta_i^I(t)\eta_i^I(t')}\equiv C^I(t-t'),\qquad R_I(t,t')\equiv R_I(t-t')$$

where $I\in\{X,Y\}$.

#### C.2 Homogeneity of strain frequencies in the $L\to\infty$, $p\to 1$, $t\to\infty$, limit

So far, we have made no assumptions and our results follow directly from the model as we have written it down. However, in order to make progress, we must we use insight from the $p = 1$ limit and make a key assumption. In this limit, all births result in mutations. While this is not necessarily a physically relevant limit, it is useful for understanding the origin of the stabilization in the stochastic model, even for $p < 1$. In the limit of $L \to \infty$ for $p = 1$, simulation evidence suggests that the strain frequencies become identical across strains (Fig. 6). So, we make the assumption that the distribution of strain frequencies at long time $t$ is approximately flat, which in turn allows us to approximate grand canonical averages (as defined above) with microcanonical averages. We check in deterministic simulations that the log strain frequencies for $p = 1$ converge to $2^{-L}$ as $L \to \infty$, as they do for weak disorder (Fig. 6). For strong disorder convergence seems to be slower, but there is still an overall trend for a fixed $\sigma$. This convergence to a uniform distribution also seems to hold for some range of $p < 1$ (that is dictated by $\sigma_a$) although the measurements are more noisy and are not presented here. We discuss the range of stability for $p < 1$ in a later section.

Note that this assumption is potentially quite strong, and primarily empirically based. However, we can take a heuristic argument for the assumption's validity in the limit. When $p = 1$, each strain will be seeded by the entirety of the interactions of its mutational neighbors and in turn seed its neighbors with the entirety of its own interactions. This will serve to average the interactions of a focal genotype over its $L$ neighboring genotypes. When $L \to \infty$, this includes infinitely many types to be averaged over, which homogenizes genotypes sufficiently well that uniformity should hold. We will see that the picture is in fact quite different for the stochastic case, where the distribution is only flat when measured over sufficiently long timescales, and that this can extend the population level stability to regimes with $p \ll 1$. This limiting homogeneity is analogous to the homogeneity of the paramagnetic phase in disordered spin models (albeit without a clear symmetry to aid in the argument). We will discuss the stochastic model in more depth as we continue.

Making the assumption that we can approximate angle bracket averages (grand canonical) with overline (microcanonical) averages in the limit, gives us the following mean-field equations for the total population size:

$$\frac{dX_{tot}}{dt} = X_{tot}\left(1 - (a+s)Y_{tot} - \sigma_a^2 \int_0^t R_Y(t-t')X_{tot}(t')dt'\right) \quad (1)$$

$$\frac{dY_{tot}}{dt} = Y_{tot}\left(-1 + aX_{tot} - \sigma_a^2 \int_0^t R_X(t-t')Y_{tot}(t')dt'\right) \quad (2)$$

We have used that the $\eta_i$ and $\eta_k$ (and the error terms $\epsilon_i$ and $\epsilon_k$) are irrelevant to the total population size in the thermodynamic limit (sums over the noise and the error end up with prefactors of $2^{-L/2}$). Note, that in a finite system, correlators of the forms $\sum_i X_i(t')X_i(t)$ and $\sum_k Y_k(t')Y_k(t)$ can potentially impact the dynamical equations through the response term. However, given that we have assumed that our system reaches a fixed point, these should be unimportant at sufficiently long times, reducing to constants that can be absorbed into the time-integrated responses (e.g. the susceptibilities). The averaged noise terms

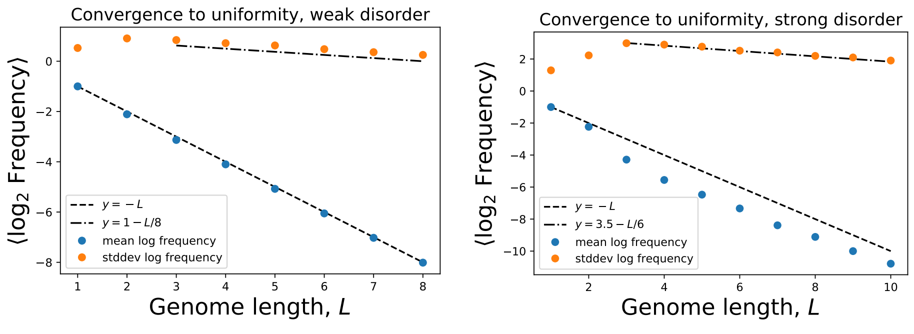


(a) (b)

Figure 6: The average $\log_2$ strain frequency converges to $-L$ for (a) weak disorder $\frac{a(a+s)}{\sigma^4} \gg 1$ (specifically, $\sigma = 1e-5$) and (b) strong disorder $\frac{a(a+s)}{\sigma^4} \ll 1$ (specifically, $\sigma = 1$). The log scale standard deviation also decays, albeit at a slower rate on this length scale, as the genome length $L$ goes to $\infty$. In both cases $p = 1$. Averages and standard deviations are calculated over 50 deterministic simulations each. Lines are to guide the eye (e.g. the decay of the standard deviation is nonlinear over longer regions of $L$). The $L = 1$ and $L = 2$ points do not fall near the line because the statistics are poor for these cases (each simulation generates only 2, and 4 strains, respectively, which do not sample enough interactions that the CLT might reasonably hold).

$\langle \eta \rangle_X$ and $\langle \eta \rangle_Y$ in turn are constant at a fixed point, and so can be absorbed by the birth and death rates.

Notice that while the invasion related noise terms are irrelevant in the limit, the feedback response terms remain relevant. This is due to the ‘strong interaction’ scaling of the disorder, e.g. $\langle \delta a_{ik}^2 \rangle = \sigma_a^2$ which should be the relevant scaling in biological systems. The disorder distribution is fixed by the set of all possible receptor-ligand pairs and is always measured in units of the birth and death rates. As such, the sum over interactions should scale with the number of interactions. There is no restriction on bacteria-phage interaction scaling in the way that there is, for instance, in the case of a many-body Hamiltonian, where the energy must be proportional to the number of particles by extensivity. Because of this, in the ‘strong interaction’ picture at the strain level, small variations in the prey growth rates and predator death rates (defined originally in the main text as $b_i$ and $d_i$ and later set to 1) can be tolerated without affecting the overall dynamics too much, i.e. interactions are dominant in determining the strain dynamics. In the ‘spin-glass’ scaling, e.g. $\langle \delta a_{ik}^2 \rangle = 2^{-L} \sigma_a^2$, assuming homogeneity, the feedback term in the total population size is also irrelevant in the $2^L \to \infty$ limit. Therefore in this picture, the total population size will follow a neutral limit cycle, and population extinction due to demographic noise or variations in the birth and death rates is a potentially important issue to address.

#### C.3 Fixed point of the DMFT equations

From the DMFT equations, we obtain fixed point equations:

$$
\begin{aligned}
0 &= 1-(a+s)Y_{tot}-\sigma^2\chi_Y X_{tot} \\
0 &= -1+aX_{tot}-\sigma^2\chi_X Y_{tot}
\end{aligned}
$$

where for clarity of presentation, we have dropped the subscript $a$, setting $\sigma_a = \sigma$. We have defined the susceptibilities:

$$\chi_X \equiv \int_0^\infty R_X(\tau)d\tau, \qquad \chi_Y \equiv \int_0^\infty R_Y(\tau)d\tau$$

We will continue to use $\sigma$ notation throughout, except when we describe the inference of the susceptibilites, where we specify $\sigma_a$ and $\sigma_s$ again for clarity's sake.

We can solve these equations to get the fixed point:

$$
\begin{aligned}
Y_{tot} &= \frac{a-\sigma^2\chi_Y}{a(a+s)+\sigma^4\chi_X\chi_Y} \\
X_{tot} &= \frac{(a+s)+\sigma^2\chi_X}{a(a+s)+\sigma^4\chi_X\chi_Y}
\end{aligned}
$$

From these expressions we can observe a few interesting things. First, in the $\sigma = 0$ case, we recover the standard fixed point for non-disordered interactions, as expected:

$$
\begin{aligned}
Y_{tot}(\sigma=0) &= 1/(a+s) \\
X_{tot}(\sigma=0) &= 1/a
\end{aligned}
$$

which is consistent with deterministic simulations, while for stochastic simulations, this regime results in extinction for explored parameter regimes.

This allows us to understand the mechanism by which the strains become homogeneous in the deterministic model. The total population size is independent of $L$, so for $L \to \infty$, $2^L$ strains have to fit into an approximately constant (at least for fixed $\sigma$) allotted abundance. In the limit, the strains have infinitesimal relative frequencies $X_i/X_{tot}$ and $Y_k/Y_{tot}$: this gives the first hint of the way to interpret the deterministic vs. stochastic dynamics. The long time frequency of the deterministic model should coincide with the long time average frequency in the stochastic model. When there is a large number of possible strains, the probability of realizing a particular one at a particular time in the stochastic dynamics is vanishingly small, whereas over longer times it is small but finite. A similar phenomenon was reported in large competitive ecological models [6], and was there referred to as 'species packing.'

However, at first glance, the $\sigma \neq 0$ expressions seem to indicate that the predator population size decreases as $\sigma$ increases. But this directly contradicts our observations from stochastic and deterministic simulations! We go on to resolve this apparent contradiction and suggest a mechanism behind the power law increase in the population size.

#### C.4 $\chi_X \chi_Y < 0$ for sufficiently large $p$

The key to understanding this discrepancy in the fixed point of the eco-evolutionary dynamics, is that for sufficiently large $p$ the susceptibilities must be such that *$\chi_Y$ is positive and $\chi_X$ is negative* when defined as we have above. This is indeed borne out in our simulations (both stochastic and deterministic, see Figs. 7 and 8), from which the susceptibilities can be numerically inferred from the *strain-level* time averages $\langle ... \rangle_t$, as was performed in [28]. [1]

Note, the inference of the response from deterministic equations is tricky over the whole range of $\sigma$ due to the presence of a crossover from mean-dominated to disorder-dominated interactions when the inverse rate $1/r_{disorder} \equiv \sqrt{\frac{a(a+s)}{\sigma^4}} \sim 1$. This crossover is discussed in depth later on and plays a crucial part in the interpretation of our observations. For the mean-dominated regime, $\sigma \lesssim [a(a+s)]^{1/4}$, the appropriate inference procedure is to work directly with the fluctuating components of the interactions to compute:

$$\sigma_a^2 \chi_X = \frac{\text{Median}_k \left[ -\sum_i \delta a_{ik} \langle X_i \rangle_t + (\sigma_a + \sigma_s) 2^{L/2} \langle \eta_k \rangle_t \right]}{2^L \text{Median}_k \left[ \langle Y_k \rangle_t \right]} \tag{3}$$

$$\sigma_a^2 \chi_Y = \frac{\text{Median}_i \left[ \sum_k (\delta a_{ik} + \delta s_{ik}) \langle Y_k \rangle_t - \sigma_a 2^{L/2} \langle \eta_i \rangle_t \right]}{2^L \text{Median}_i \left[ \langle X_i \rangle_t \right]} \tag{4}$$

where we have temporarily reintroduced $\sigma_a$ and $\sigma_s$ for clarity. By taking the time averages first, then taking the medians over $k$ and $i$, we remove the noise terms (which have zero mean) and get the susceptibilites $\chi_X$ and $\chi_Y$ in terms of measurable quantities. These can then be averaged over simulations.

However, in the strong disorder regime, $\sigma \gtrsim [a(a+s)]^{1/4}$, the appropriate inference procedure is to take:

$$\sigma_a^2 \chi_X = \frac{\text{Median}_k \left[ -\sum_i a_{ik} \langle X_i \rangle_t + a \langle X_{tot} \rangle_t + (\sigma_a + \sigma_s) 2^{L/2} \langle \eta_k \rangle_t \right]}{2^L \text{Median}_k \left[ \langle Y_k \rangle_t \right]} \tag{5}$$

$$\sigma_a^2 \chi_Y = \frac{\text{Median}_i \left[ \sum_k (a_{ik} + s_{ik}) \langle Y_k \rangle_t - (a+s) \langle Y_{tot} \rangle_t - \sigma_a 2^{L/2} \langle \eta_i \rangle_t \right]}{2^L \text{Median}_i \left[ \langle X_i \rangle_t \right]} \tag{6}$$

This inference is more appropriate since the fluctuating part of the interactions is comparable to the mean in this regime due to the power law growth of the total population sizes. Therefore both terms must be accounted before taking the median. However, this inference is also limited due to potentially slow convergence of the deterministic simulations. Despite improvements in the inference of the response using this method (see Fig. 10), it was still difficult to obtain numerically accurate enough estimates to get quantitative measurements in the strong disorder regime. We will discuss this regime in more depth shortly.

[1] Note that the change of sign of carrying capacity due (separately) to quenched disorder, diffusion and demographic noise has been reported in the context of a generic model of directed percolation [17]. Since all three are at play in the full stochastic model and strong diffusion (mutation) occurs even at the deterministic level, we expect that a similar effect is at play here. Care would need to be taken in checking this and a full renormalization group analysis of the strain level equations would need to be carried out to verify this analytically.

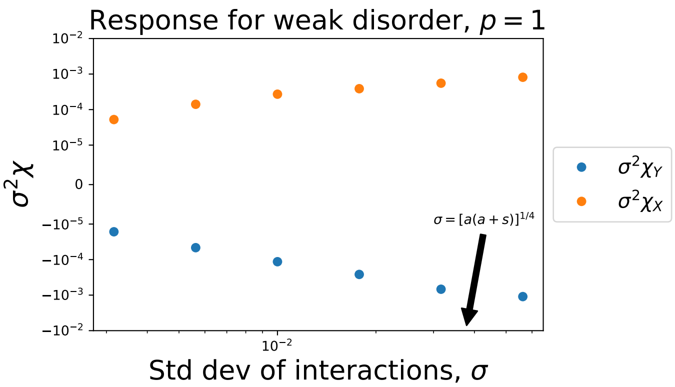


Figure 7: Inferred predator and prey response functions for deterministic simulations with $L = 8$ in the weak disorder regime, $\sigma \lesssim [a(a+s)]^{1/4}$. Each point is the median over 20 simulations. As conjectured, $\chi_Y < 0$ and $\chi_X > 0$, over the entire range of $\sigma$ simulated. In the mean-dominant regime, the product of the inferred response terms is smaller in magnitude than $a(a+s)$, and the magnitudes of the responses themselves ($\chi_X$ and $|\chi_Y|$) are approximately independent of $\sigma$.

The result $\chi_Y < 0$ can be understood intuitively to be a result of mutation, which changes the nature of the feedback. Mutation has not entered into the total population size dynamics up to this point although it crucially enters into the individual strain dynamics (which enter into the susceptibility measurement in the first place). We can get a handle on this intuition in the strong mutation regime ($p \sim 1$). In this regime, some large fraction of predator births (which only occur due to predation interactions) result in random mutation. So when predator $Y_k$'s favorite prey $X_i$ is at high abundance, much of $Y_k$'s consumption results in growth of its mutational neighbors. These mutational neighbors in turn are unlikely to prefer the same prey (if the interactions are sufficiently uncorrelated). So at the end of the feedback loop $X_i$ gets a small boost in growth rate since overall its' strongest predator $Y_k$ has been relatively depleted due to its constant death rate. Prey mutations do not feed back on the response $\chi_X$ since they do not depend on the interactions. [2] See Fig. 9 for a schematic

[2]N.B. In similar models with *mutation rates*, $\mu$, which exhibit regular diffusion in genotype space (versus *mutation probabilities*, $p$, which exhibit nonlinear diffusion, see below strain level DMFT equations), there should also be a stable ergodic phase but only if $\mu$ is sufficiently large. More specifically the important ingredient is that the predator-specific $\mu$ is sufficiently large (i.e. $L\mu^{-1}$ should be comparable to the timescale of the interactions) that it makes $\chi_Y$ negative. In phage-bacteria systems this may in fact be the case since phage and viruses generally have significantly higher mutation rates than microbes. In the end we argue that the $p$-model is more true to the underlying biology than the $\mu$-model. However, the $\mu$-model might be more readily compared to existing data, since what is typically measured is a rate rather than a probability. A more direct and detailed comparison of the two models is an interesting avenue for future work but is beyond the scope of the present study.

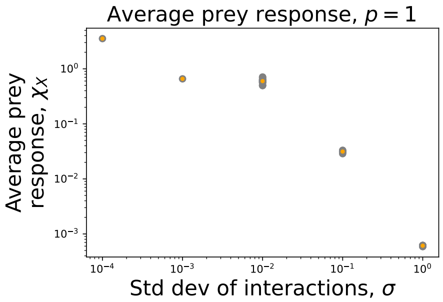


(a)

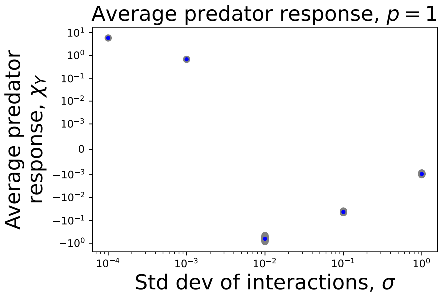


(b)

Figure 8: Average inferred response over a range of $\sigma$ for $p = 1$ in stochastic simulations (using the strong disorder method), initialized with a uniform distribution of predator and prey. Gray dots are individual simulations, colored dots are averages over simulations. (a) Prey response $\chi_X$ is positive over the whole range. (b) Predator response $\chi_Y$ plotted on a symlog scale. Note the positive response to the left of the panel (b) is due to the inference method. The sum of the different terms in the numerator of Eqs. 5 and 6 is unbalanced, which distorts the computation of the median, which is a nonlinear operation. In this case, $(a + s)\langle Y_{tot}\rangle_t$ is only sufficiently large to contribute in the regime $\sigma^2 \gg \sqrt{a(a + s)}$. However, using the weak disorder response inference demonstrates a negative $\chi_Y$ with $|\chi_Y \chi_X| \sim \mathcal{O}(1)$, for a finite fraction of simulations with these values of $\sigma$, since they fall close to either side of the boundary of the stable regime (as shown in Fig. 3 of the Main Text).

illustration of this.

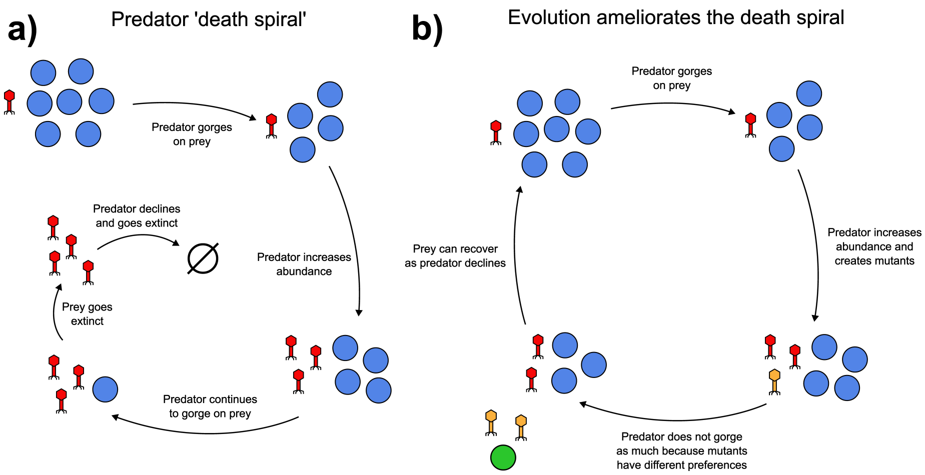


Figure 9: Schematic of the types of feedback on a focal prey strain both a.) without evolution (negative) and b.) with evolution (positive).

#### C.5 A crossover from weak to strong disorder suggests the origin of the population size power laws

Once we see that $\chi_X \chi_Y < 0$, we can also begin to understand why the population size increases with $\sigma$ from self-consistency requirements. For the fixed point expressions to be valid, they are required to be positive. We see for fixed large but finite $2^L$, as we increase $\sigma$, the susceptibilities $\chi_Y$ and $\chi_X$ must become sufficiently small in compensation such that the following constraint is satisfied for the fixed point equations to remain valid:

$$\sigma^4 |\chi_X \chi_Y| < a(a+s)$$

The $\chi_i$ might become small since the relevant rate variables are positive random variables. When drawn from a distribution with high variance (but positive support), many will be small in magnitude and some will be large. Therefore, the $\chi_i$ should not become arbitrarily small since at least some of the interactions are quite large in the intermediate regime $2^{2L} \gg \sigma^4 \gg a(a+s)$ where linear response is still a valid approximation. This constraint reveals a crossover from mean-dominated to disorder-dominated behavior when the inverse rate $1/r_{disorder} \equiv \sqrt{\frac{a(a+s)}{\sigma^4}} \sim 1$. This rate is what sets the chaotic fluctuations due to disorder and is intimately related to the May transition [18, 4].

In the small $\sigma$ regime, the products $\sigma^2 \chi_X$ and $\sigma^2 \chi_Y$ should be small since the dynamics are mean-dominated (e.g. the equilibrium values $X_{tot}$ and $Y_{tot}$ will be approximately independent of $\sigma$, which holds in the small $\sigma$ regime of

Fig. 3). We make the scaling ansatz for the asymptotic expansion of the product of the susceptibilities as $\sigma$ becomes large (the disorder-dominated regime):

$$\sigma^4|\chi_X\chi_Y| \quad \sim \quad a(a+s) - c\left(\frac{p}{\sigma}\right)^\eta$$

where $c$ is a constant that gives the correct dimensional units and the quantity in brackets is an expansion parameter. This scaling form can be thought of as a bound on how large the magnitude of the susceptibilities can be and still allow for a valid nontrivial fixed point.

In the mean-dominated regime, we in fact see the expected approximate $\sigma$-independence in the population size in deterministic simulations (see Fig. 3). For the disorder-dominated regime, we were unable to directly uncover the hypothesized scaling due to problems with convergence of the deterministic simulations, which in turn made it difficult to infer the response. While this could also potentially indicate a breakdown of the fixed point assumption (akin to the 'multiple attractors' phase identified in [4, 31]), we see no compelling evidence of this in the $p = 1$ simulations (e.g. there are no strong, persistent fluctuations). However, for low $p$, there can be persistent strong fluctuations for a sufficiently large $\sigma$. We discuss aspects of this shortly, especially how it might relate to the stochastic simulations. However, we save an in-depth study for future work.

The key difficulty with the strong disorder regime is that the disorder and the mean interactions are of comparable magnitude, so they need to be treated simultaneously. This amplifies any transient fluctuations in the strain dynamics when inferring the response. While we cannot infer the scaling of the second term of the expansion quantitatively given our computational constraints, we extract the correct first order scaling since the inferred response is $\mathcal{O}(a(a+s))$. See Fig. 10 for $\sigma^4|\chi_X\chi_Y|$ inferred from deterministic simulations for $p = 1$.

Despite the issues with the numerical accuracy of the deterministic simulations, we see evidence of the conjectured scaling in the stochastic simulations at large $\sigma$ (see Fig. 11). Once again, this evidence is quantitative only to leading order since the statistics are too poor to extract the exponent of the second term of the expansion directly. However, qualitatively, aspects of the scaling ansatz agree with the inferred responses, even for $p \ll 1$. Moreover, the conjectured scaling is made apparent in the population sizes in both the stochastic and deterministic simulations.

Specifically, as we increase $\sigma$ beyond the crossover, we expect the population size to diverge:

$$X_{tot} \sim Y_{tot} \sim \sigma^\eta$$

This is borne out in our stochastic simulations with a value for the exponent $\eta \approx 1/4$ and in the above deterministic simulations $\eta \approx 1/2$. Furthermore, we expect the fixed point population size to decline with $p$:

$$X_{tot} \sim Y_{tot} \sim p^{-\eta}$$

which also agrees with stochastic simulations. For deterministic simulations, a finite range of $p < 1$ seems to result in a fixed point at long times. Below

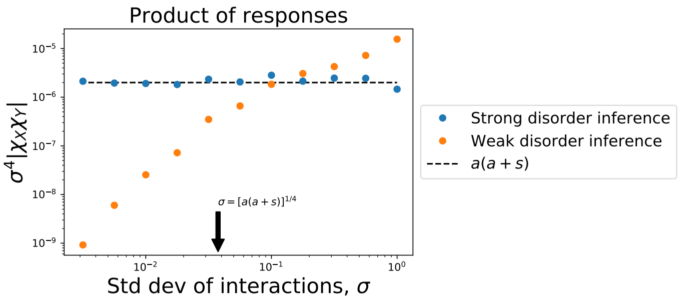


Figure 10: Absolute value of the product of the response functions $\sigma^4|\chi_X\chi_Y|$, inferred from the strain dynamics in deterministic simulations with $L = 8$, $p = 1$. Points are the median over 20 simulations. Near the crossover, the product $\sigma^4|\chi_X\chi_Y|$ approaches $a(a+s)$. However, above the crossover, the weak disorder inference procedure breaks down. While the strong disorder inference procedure infers a quantity that remains $\mathcal{O}(a(a+s))$, the measurements are still too noisy to extract quantitative information about the singular term of the expansion in this regime.

this range we observe what are potentially chaotic dynamics. For deterministic simulations at fixed $\sigma$ and $L$, the viable range of $p$ can be relatively small (as well as dependent on $\sigma$), as discussed in a later section. However, stability is observed over a much broader range of $p$ in the stochastic model.

Finally, we note that while the mean of the anti-symmetry breaking interactions, $s$, can affect the location of the fixed point, the variance $\sigma_s$ is irrelevant. These interactions do not form a closed loop between predator and prey, so there is no feedback, which is how variability comes in at the population level. However, the variability of these interactions can contribute to the stochastic dynamics. In addition the $s_{ik}$ (and $\sigma_s$) will certainly be relevant in the short time strain level dynamics as well as the details of the time-dependence of the response.

##### C.6 Stability of the mean field fixed point

We can perform a linear stability analysis of the equations:

$$\frac{dX_{tot}}{dt} = X_{tot}\left(1 - (a+s)Y_{tot} - \sigma^2 \int_0^t R_Y(t-t')X_{tot}(t')dt'\right)$$

$$\frac{dY_{tot}}{dt} = Y_{tot}\left(-1 + aX_{tot} - \sigma^2 \int_0^t R_X(t-t')Y_{tot}(t')dt'\right)$$

We focus on $\omega = 0$ mode which is maximally unstable, and which simplifies our analysis since this allows us to set the responses to their $\tau = \infty$ values, $\chi_X$ and

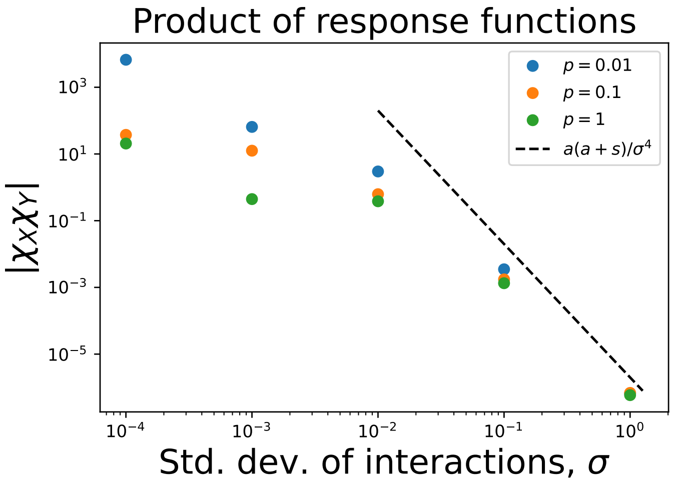


Figure 11: Absolute value of the product of the inferred response functions $|\chi_X \chi_Y|$, averaged over stochastic simulations (inferred with the strong disorder method), initialized with a uniform distribution of predator and prey. To leading order the product scales with $\sigma^{-4}$, getting closer to $\frac{a(a+s)}{\sigma^4}$ as $\sigma$ increases, as predicted (with no fitting parameters). Surprisingly, the scaling ansatz works for $p \ll 1$, suggesting that the stabilization mechanism is valid over a range of $p$. Furthermore, points with higher $p$ are farther from the dashed line, as qualitatively predicted by the theory. Accurately measuring the scaling of the second term (the deviation from $a(a+s)/\sigma^4$) was not feasible due to a lack of sufficient numerical precision. However, this scaling is made apparent in the population size.

$\chi_Y$ [24]. A simple calculation leads us to the stability eigenvalues:

$$\lambda_{\pm} = \frac{-1 \pm \sqrt{1 - 4\frac{[(a+s)+\sigma^2\chi_X][a+\sigma^2|\chi_Y|]}{a(a+s)-\sigma^4|\chi_X\chi_Y|}}}{2}$$

where we have the discriminant:

$$\frac{\left[(a+s)+\sigma^2\chi_X\right]\left[a+\sigma^2|\chi_Y|\right]}{a(a+s)-\sigma^4|\chi_X\chi_Y|} > 0$$

since, importantly, the denominator must be greater than 0 for the sake of self-consistency of the fixed point equations. Therefore the total population size of a collection of predator-prey pairs is a stable focus although the dynamics of a single predator-prey pair is a neutrally stable limit cycle. In the limit of strong disorder, the complex component of the stability eigenvalue grows so that the frequency of the oscillatory component increases, potentially interfering with residual correlation timescales or response timescales in finite systems (which could cause the fixed point assumption to potentially breakdown for $p < 1$, which we discuss imminently). Interestingly, for weak disorder (e.g. below the crossover scale) we have that the mean field population size fixed point is a focus with eigenvalues:

$$\lambda_{\pm} = \frac{-1 \pm \sqrt{3}i}{2} + \mathcal{O}(\sigma)$$

whereas the non-zero fixed point of the equivalent neutral predator prey model has pure imaginary stability eigenvalues:

$$\lambda_{\pm}^{neutral} = \pm i$$

so that disorder with evolution is a singular perturbation at the deterministic level. This is borne out is deterministic simulations, where the dynamics of the total population size seem to be a stable focus for small $\sigma$ (see Fig. 12). We discuss situations with $p < 1$ in the next section.

Despite the deterministic stability, there are potentially large transient initial fluctuations and also metastable states which are observed in both deterministic and stochastic simulations. Furthermore, in the stochastic simulations, we see that oscillatory fluctuations persist. We conjecture that this is a form of stochastic resonance in which the stochasticity (demographic noise) has frequency components that are comparable to the frequency of the damped oscillations at the mean field level.

#### C.7 Summary of the DMFT analysis and large $\sigma$, finite $p < 1$

In this analysis, we have identified several interesting phenomena having to do with the deterministic equations describing predator-prey co-evolution. First, we have shown that the strains become more and more homogeneous as the genome length $L \to \infty$, suggesting that in the limit, the distribution of strain frequencies is uniform. Then, we have employed this fact to show that the total population size of the predator and prey populations can reach a steady state for sufficiently high $p$. With an examination of the steady state expressions, and measurements from simulations, we have shown that the predator response $\chi_Y$ is negative. There is a crossover in the behavior of the response functions, when the system goes from a 'mean-dominated' regime (the inverse rate $\sqrt{a(a+s)}/\sigma^2$ is large) to a 'disorder-dominated' regime (the inverse rate $\sqrt{a(a+s)}/\sigma^2$ is small). In the disorder-dominated regime, we see a power law increase in the population size of both the predator and the prey types. Importantly, we see consistency between the theory corresponding to the deterministic dynamics and simulations of the stochastic dynamics. Finally, we have shown that the predator and prey population fixed points are stable in the $L \to \infty$ limit, with theory and simulation suggesting stability for broad range of $\sigma$, see Fig. 12. Whether this stability extends down to $p \to 0^+$, is an important topic to explore, although the fact that $p$ did not enter into the stability calculations (through $\chi_Y$ or $\chi_X$) for $\sigma \to 0^+$ suggests that it should for sufficiently weak disorder.

For fixed finite $\sigma$, as $p$ is decreased, there seems to be a $\sigma$ dependent transition from stability to a persistently chaotic state. This can be understood as a transition that occurs when the mutation timescale is comparable to an appropriate timescale set by the disorder. The longer of the two mutation timescales is given by the prey mutation term (the predator mutation term is $\gtrsim$ the prey mutation term), so we take the prey mutation timescale:

$$t^X_{mut} = \frac{1}{pb_i} = \frac{1}{p}$$

where we have used that the birth rates are all uniform, $b_i = 1$. In order for the fixed point solution to remain self-consistent, mutation should act more quickly than any effects due to interactions (mutation should homogenize strain abundances before disorder can disrupt them). We can now compare this timescale to an appropriate 'disorder timescale.' We can estimate this by taking the inverse of the disorder rate:

$$t^X_{disorder} = const. \times \frac{\sqrt{a(a+s)}}{\sigma^2}$$

where the constant is make sure the units are correct. This timescale need not be particularly accurate (in terms of the constant numerical prefactor) since we are interested in rough qualitative behavior rather than in the precise location of this transition. Rather, we assert intuitive requirements for the dependence on $p$ and $\sigma$. The disorder timescale gets shorter with increasing $\sigma$, e.g. more disordered systems are more likely to exhibit chaotic behavior on short times (which is consistent with simulations). The magnitude of the disorder timescale should not diverge as $p \to 0$. It should rather saturate to some finite quantity or potentially go to zero when $p = 0$. It should only potentially diverge when $p \to 1$. Therefore the disorder timescale can only be weakly dependent on $p$, with at most a leading order dependence $1/(1-p)$, which will not affect our qualitative results. We do not explicitly assume this sort of divergence at $p = 1$ to be as stringent as possible in our comparison of the magnitudes of mutation and disorder. We look to simulations to confirm our qualitative predictions afterwards.

So roughly we must have:

$$t_{mut} \lesssim t_{disorder}$$

for stability. In the weak mutation regime, this is achievable for any $\sigma$ so long as $p$ is sufficiently large, over an $\mathcal{O}(1)$ range (the timescale $\frac{\sqrt{a(a+s)}}{\sigma^2}$ is greater than one over this regime). However, in the strong disorder regime, we have:

$$const. \times \frac{\sigma^2}{p} \lesssim 1$$

In this regime, for large $\sigma$ only a potentially small range of mutation probabilities $1-p_{min} \sim \mathcal{O}(1)$ can rescue stability (the left hand side diverges as $p \to 0$). When this happens, mixing due to mutation does not occur on a timescale fast enough to suppress the disorder effects, which drive chaotic fluctuations (whereas in the $p = 1$ case, while there are still interactions, they all go towards mixing through mutation, and stability persists for all ranges of $\sigma$). Therefore for fixed $\sigma$ and low enough $p$, the fixed point solution breaks down.[3] We demonstrate this in Fig. 13. The timescale argument allows us to understand why the deterministic simulations do not seem to have stability extend down to $p \ll 1$. However it leaves a gap in our understanding of the stochastic simulations, where stability seems to extend up to large $\sigma$ for a broad range of $p$. How can the stochastic simulations achieve apparent population stability (when looking on the mean over a time window) in a parameter regime where the deterministic simulations are observed to be unstable?

The solution to this apparent mystery is that demographic noise plays a key role in stabilizing the predator-prey system. Demographic noise sets a 'lattice spacing' so that, at a given time, the system does not 'see' all the strains (e.g. many strains are extinct at a single time). Therefore, in the stochastic model, the relevant comparison is between the mutation timescale and an *effective* disorder timescale defined by the strains that are present at a given time $t$. This effective disorder timescale (which has been regularized by demographic noise) may be significantly greater than the 'full' disorder timescale since, though all strains are present when observing over very long times, any particular strain is likely extinct at a given time. In addition, demographic noise (and resultant extinction) more efficiently homogenizes abundances as averaged over a fixed time, further 'flattening' long time strain abundances, so that we expect aspects of the mean-field results to hold (e.g. the long time response functions) when observing the stochastic system over long times. We discuss the details of the stochastic model in the next section.

[3]This is a version of the classic May result [18] and the results in [4, 30]. When $\sigma^2$ gets too large, the fixed point solution breaks down for $p < 1$.

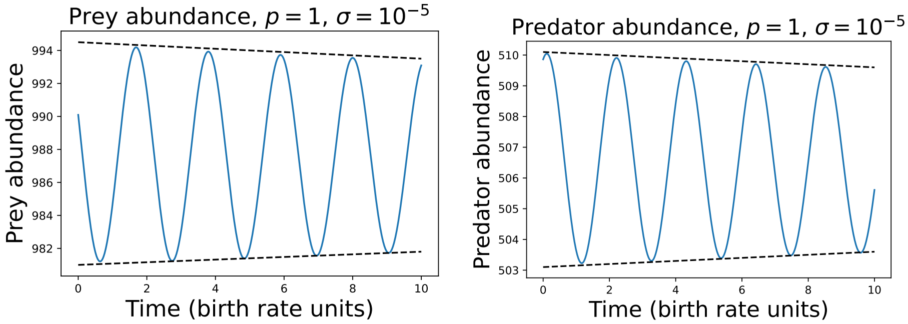


(a) (b)

Figure 12: (a) Prey and (b) predator abundances for low $\sigma$, $L = 8$. The oscillation amplitude decreases over time, in contrast to the predator-prey model with $\sigma = 0$ (neutrally stable limit cycle). Dashed lines are to guide the eye. Note the range on the y-axis is quite small. Despite the evidence for stability at low $\sigma$ on a short time scale, because of difficulties in simulating trajectories to sufficient numerical accuracy, longer timescales were difficult to achieve.

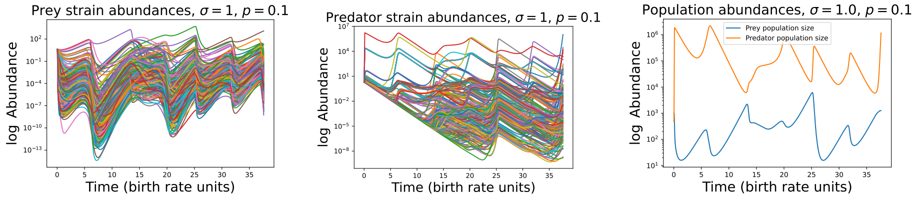


(a) (b) (c)

Figure 13: (a) Prey strain, (b) predator strain and (c) total population abundances in the mean-field for $p = 0.1$, $\sigma = 1$, $L = 8$. The dynamics are unstable if there is no possibility of extinction. The population dynamics also appear chaotic in this regime. For $p = 0.1$, $\sigma = 1$ the stochastic dynamics exhibit apparent stability.

### Appendix D Stochastic model

#### D.1 The effective disorder, $\sigma_{eff}$

Due to strain extinctions, the sample of active strains can have a potentially very different distribution of interaction rates than the full set of possible interaction rates. Moreover, this quantity dynamically adjusts itself as the dynamics proceed, when new strains are seeded and older strains go extinct. If we look at the trajectory of a stochastic simulation, we see that the effective disorder $\sigma_{eff}$ can be much lower than the true $\sigma$, although it can fluctuate over several orders of magnitude. By self-organizing to a state with lower disorder at a given time (on the median), the stochastic dynamics are able to circumvent the issue with instability in the strong disorder regime of the deterministic model. However, long-time average quantities, like the susceptibility, 'see' the full disorder since the population traverses the genotype space and achieves a nearly uniform distribution at (potentially very) long times. In this way, the stochastic model is still able to realize the increase in total population size observed in the deterministic model at large $\sigma$, albeit with a different exponent, while maintaining population-level stability over a broad range of $p$. We conjecture how this exponent might be generated in a later section.

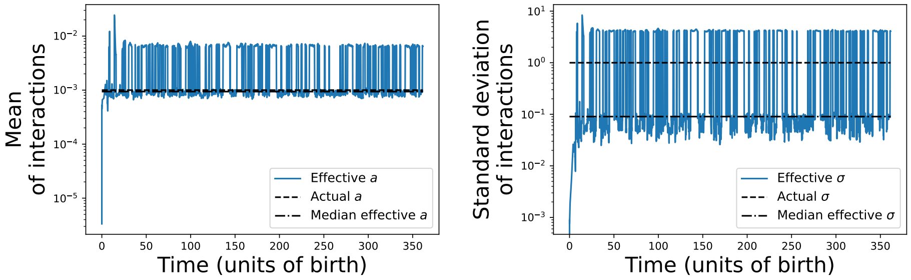


(a) (b)

Figure 14: (a) Effective $a$ and (b) effective $\sigma$ for $a = 0.001$ and $\sigma = 1.0$, $p = 1$. The realized effective $a$ of active strains is of the same order as its actual value. We see that the realized effective $\sigma$ at a given time can be much lower than its full value. On the median, it is much lower than its full value, though it fluctuates quite rapidly between extremes on a log scale. Because the effective $\sigma$ is small, this allows the total population size to avoid chaotic dynamics that might be present for these parameters in the deterministic model.

We can now run the timescale comparison for the stochastic model, where we have the mutation timescale:

$$t^X_{mut} = \frac{1}{p}$$

However, the correct disorder timescale depends on what is observed *locally* – only strains that are present at a given time will be able to drive chaotic fluctuations. This is given by:

$$t_{disorder}^{eff} = const. \times \frac{\sqrt{a(a+s)}}{\sigma_{eff}^2}$$

where we assume the $\sigma_{eff}$ can depend on $p$, $\sigma$, $a$, and $s$ in unknown ways. We can understand intuitively that $\sigma_{eff}$ that $\sigma_{eff}$ gets smaller as $p$ gets smaller (if mutations arrive less frequently, there will be less diversity typically). This dependence on $p$ mitigates the divergence as $p \to 0$ for large $\sigma$ in the stability constraint:

$$const. \times \frac{\sigma_{eff}^2}{p} \lesssim 1$$

This allows that a long time homogeneous fixed point can be valid so long as the predator doesn't stochastically go extinct due to demographic noise, which we discuss in the next section. One subtle observation in the stochastic simulations is that while demographic noise mitigates the destabilizing effect of the strong disorder regime, the power law scaling seems to extend over a much broader observable range than in the deterministic model (e.g. into the weak disorder regime). A more detailed analysis of the stochastic dynamics might uncover the precise (dynamical) mechanism by which this comes about.

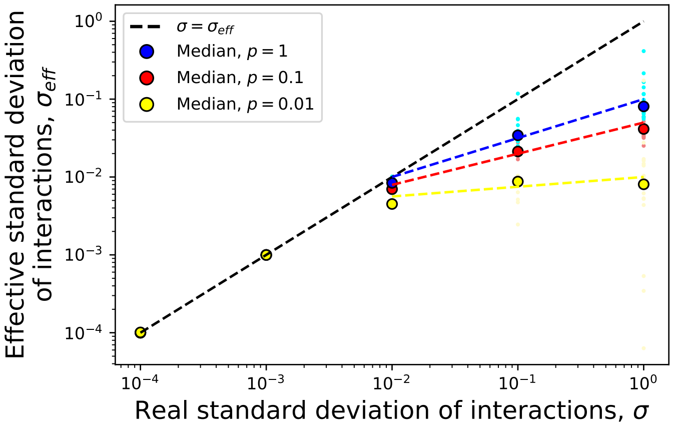


Figure 15: Median $\sigma_{eff}$ for a range of $p$ and $\sigma$. Small dots correspond to $\sigma_{eff}$ for individual simulations. Large dots are medians over all simulations. Colored dashed lines are to guide the eye.

#### D.2 Heuristic analysis of predator extinction due to stochasticity

Any rigorous analysis of the full stochastic dynamics is bound to be extremely difficult. The complicated form of the strain-level reaction-diffusion equations (which at the DMFT level is a nonlinear diffusion with unknown memory kernels) resists obvious solution. However, we are equipped to heuristically discuss these dynamics, choosing to specifically focus on the location of the transition from a 'phase' where the predator population goes extinct on a timescale comparable to the neutral model ($\sigma = 0$) to a 'phase' where predator extinction happens on a much longer timescale due to the stabilizing effects of mutation. We refer to the former as the 'extinct' phase. We refer to the latter as the 'stable' phase. However, these terms are used non-rigorously since extinction is inevitable for any finite stochastic system with an absorbing state.

In the predator extinction process, there are three timescales that are important to compare. First is the extinction timescale $t^Y_{ext}$, which is the typical time that the predator population will take to go extinct without adaptation. Second is the timescale for the *population* to produce a single mutant, $t^Y_{mut}$. Third is the diffusion timescale, $t^Y_{diff}$, which is the timescale that a *single individual* produces a beneficial mutant. By definition, $t^Y_{diff} \gg t^Y_{mut}$, by a factor of the order the total population size. We expect that these timescales differ depending on the direction we approach the transition point, and with which control parameter we use to do so, $\sigma$ or $p$.

We start by analyzing the transition in $\sigma$, which is the simpler case. For this scenario, it will be more informative to start in the stable phase and lower down to the extinct phase. We start with the diffusion timescale, since it is the longer timescale and will be the first to cross the extinction timescale. The mutation term for the $k^{\text{th}}$ predator strain can be written:

$$p \sum_{l \in \mathcal{S}} \frac{1}{L} \left( \sum_i a_{il} X_i Y_l - \sum_i a_{ik} X_i Y_k \right) \tag{7}$$

so that we can use DMFT to get:

$$\begin{aligned} p \sum_{l \in \mathcal{S}} \frac{1}{L} \Bigg[ & \left( a X_{tot} + \sigma 2^{L/2} \eta^X_l - \sigma^2 2^L \int_0^t R_X(t-t') Y_l(t') dt' \right) Y_l \\ & - \left( a X_{tot} + \sigma 2^{L/2} \eta^X_k - \sigma^2 2^L \int_0^t R_X(t-t') Y_k(t') dt' \right) Y_k \Bigg] \end{aligned}$$

Then, employing the nearly uniform limit for $t \to \infty$ (which relies on summing $L$ terms instead of $2^L$ as in Supplement C), we have the time-averaged mutation term:

$$p \left( a X_{tot} - \sigma^2 \chi_X Y_{tot} \right) \sum_{l \in \mathcal{S}} \frac{Y_l - Y_k}{L}$$

The sum over differences of abundances is the Laplacian operator on the hypercube. The expression in front is an effective diffusion coefficient, which is approximately constant in the stable phase since $X_{tot}$ and $Y_{tot}$ are approximately constant. We have that the diffusion timescale is simply:

$$t_{diff}^Y = \frac{l_D^2}{\langle D_Y \rangle}$$

where the averaged diffusion constant can be approximated by $\langle D_Y \rangle \approx aX_{tot} - \sigma^2 \chi_X Y_{tot}$ in the thermodynamic limit.

The distance $l_D$ can be understood as how far in genotype space a focal type $Y_k$ needs to mutate to reach a beneficial genotype. Typically there is a dimension dependent factor connecting the mean squared distance to a diffusion time. However given that the interactions are randomly drawn, we can approximate that roughly half will be beneficial and half will be deleterious. Therefore a randomly mutating predator will need to undergo much fewer than $L$ 'trials' before finding a beneficial mutation with high probability. In the thermodynamic limit $L \to \infty$ the distance $l_D$ is $\mathcal{O}(1)$. We set $l_D = 1$ for clarity of exposition.

Therefore, the diffusion timescale is given (in the scaling limit) by:

$$t_{diff}^Y = \frac{1}{p\left(aX_{tot} - \sigma^2 \chi_X Y_{tot}\right)} \approx \frac{1}{paX_{tot}}$$

where the last approximation holds for sufficiently low $\sigma$. Furthermore, it can be established that:

$$t_{ext}^Y \sim Y_{tot}$$

for a neutral limit cycle subject to demographic fluctuations [25, 26, 27, 2] – this results from diffusive population size fluctuations. This should be the correct scaling of the extinction timescale for low enough $\sigma$, since all strains are approximately neutral with respect to each other. As $\sigma$ increases, this timescale will increase continuously.

Therefore we have the ratio of timescales:

$$\frac{t_{ext}^Y}{t_{diff}^Y} = paX_{tot}Y_{tot} \approx pac_1 \left(\frac{\sigma}{p}\right)^{\eta} c_2 \left(\frac{\sigma}{p}\right)^{\eta}$$

where we have used that the population sizes in the stable phase obey power laws in $\sigma/p$, which gives us the right $\sigma$-dependence. We see that when this ratio is $\gtrsim 1$, we are in the stable regime as shown in Fig. 3a in the Main Text.

We also see that there is some persistence for $\sigma < \sigma_{diff}$, where $\sigma_{diff}$ inferred using the condition $\frac{t_{ext}^Y}{t_{diff}^Y} \sim 1$. Due to finite size effects, finite simulation times and particular realizations of disorder, individual simulation trajectories can persist over a fixed window for $\frac{t_{ext}^Y}{t_{diff}^Y} \lesssim 1$. This 'smearing' is due to the shorter mutation timescale $t_{mut}^Y$, which can roughly be estimated with the integral condition:

$$\int_0^{t_{mut}^Y} paX_{tot}Y_{tot}dt \sim 1$$

so that:

$$t_{mut}^Y \sim \frac{1}{paX_{tot}Y_{tot}dt}$$

. For $t^Y_{ext}/t^Y_{mut} \lesssim 1$, the predator population will almost certainly go extinct on the observed timescale.

We now consider holding $\sigma$ fixed and varying $p$ by coming from $p = 0$, e.g. from the extinct phase to the stable phase. We choose this direction since we typically consider the situation where we initialize with a single pair of strains and wait for them to diversify. When starting with a single strain the extinction vs. non-extinction distinction is relatively simple: a strain with net positive growth rate must evolve before the first strain has time to go extinct. We first consider the shorter timescale, $t^Y_{mut}$, the timescale that the *population* creates a beneficial mutant.

The population mutation timescale (as averaged over a window) can be given by the condition that $\mathcal{O}(1)$ mutants are produced over $t^Y_{mut}$, which is expressed as following integral condition:

$$\int_0^{t^Y_{mut}} paX_{tot}Y_{tot}dt \sim 1$$

Using the mean value theorem to replace $X_{tot}$ and $Y_{tot}$ (which are generally time dependent) with their mean values and solving for $t^Y_{mut}$, we get:

$$t^Y_{mut} \sim \frac{1}{pY_{tot}}$$

Comparing this to the extinction timescale $t_{ext} \sim Y_{tot}$, we get:

$$\frac{t^Y_{ext}}{t^Y_{mut}} \sim pY^2_{tot} \approx \frac{p}{(a+s)^2}$$

In the region where $p$ is larger than $(a+s)^2$ but not too large, the population survives chance mutation by chance mutation, while consisting of $\mathcal{O}(1)$ strains at a given time (see Fig. 3c in the Main Text for confirmation in simulations). This regime is hypothesized to be 'metastable' – with a potentially longer extinction time than that in the neutral limit, but not necessarily significantly longer.

However, as we continue to increase $p$, the diffusion timescale (the timescale that a single individual will seed a mutation) becomes more and more relevant when compared to the extinction time, which remains fixed on average since the population only consists of a single pair of strains at a time. Given the diffusion process in genotype space, as before, we have:

$$t^Y_{diff} = \frac{l_D^2}{\langle D_Y \rangle}$$

The averaged diffusion constant can be approximated by $\langle D_Y \rangle \approx pa\langle X_{tot} \rangle_t$ where $\langle X_{tot} \rangle_t$ is the time-averaged abundance of the average prey strain. When there are multiple strains present, this may not be a good approximation, but since we start in the metastable regime, where there are only $\mathcal{O}(1)$ active strains at a time, this should be a reasonable approximation to leading order. We have the same extinction timescale as before (since we start with a single strain):

$$t^Y_{ext} \sim Y_{tot}$$

Taking the ratio of the extinction and diffusion timescales gives the ratio:

$$\frac{t^Y_{ext}}{t^Y_{diff}} \equiv paX_{tot}Y_{tot} \approx \frac{p}{a+s}$$

where $a\langle X_{tot}\rangle_t \approx 1$, e.g. conditioning on non-extinction, the mean prey abundance times the interaction rate should be $\approx 1$ to balance the predator death rate.

This quantity is required to be $\gtrsim 1$ for multiple strains to circulate at a time. This is consistent with Fig. 3c in the Main Text, since the strain entropy at $t_{final}$ is approximately zero (on the median) below this point. We notice that the population size scaling only starts immediately above $p^* = a + s$. For the region $(a + s)^2 < p < a + s$, there are potentially long-lived transients due to initialization or a particular realization of the random interactions. Such putative metastable scenarios are potentially interesting to study in more depth and are potentially relevant to understand the process of colonization by random strains in experimental and natural systems. However, a detailed exposition of metastability is beyond the scope of the present study.

We expect that the diffusion and extinction timescales will differ as we approach the transition from both directions varying $p$ (from the metastable regime vs from the stable phase), so that the transition between the metastable phase and stable phase will be first order when $p$ is used as a control parameter [11]. This is due to the fact that approaching the transition point from the metastable phase, depending on the initial conditions, we move along a neutrally stable limit cycle or divergent chaos (both of which rapidly wander or decay to extinction), where the typical population sizes are set by $X_{tot} \sim 1/a$ and $Y_{tot} \sim 1/(a + s)$, and so the ratio $t_{ext}/t_{diff}$ only depends on $p$ linearly. Whereas to the right side of the transition the total population sizes are substantially larger because they follow the power law scaling relation (see Fig. 3 in the Main text – the median population size increases as $p$ is decreased from 1, until it drops off past the transition), so that simultaneously $t_{diff}$ gets smaller and $t_{ext}$ gets bigger as the transition is approached from $p = 1$.[4] Therefore we predict that predator-prey eco-evolutionary dynamics should exhibit a form of hysteresis in the mutation rate, whereby an initially stable population can quasi-statically have its mutation probability lowered beyond the transition point into the metastable regime and remain stable for some range of $p$. However, when using $\sigma$ as the relevant

[4] N.B. We can make an alternate argument for why the transition should be 1st order. Grassberger and Janssen conjectured that any active-inactive state transition should fall under the directed percolation (DP) universality class [13, 14]. Our transition is an active-inactive state transition so it should fall under DP. In $\sigma$, we expect that the transition is second order. In $p$, we expect that the transition is first order. Above the critical dimension, there can exist a 1st order transition [13] (when the carrying capacity is the control parameter, as is effectively the case in our model, since $p$ flips its sign). Our model ($d = 2^L$) is well above $d_c = 4$ so this is still consistent with a first order transition. It is interesting to note that the single locus dynamics ($L = 1$) are below the critical dimension and the two-locus dynamics ($L = 2$) are at the critical dimension. So the transition in $p$ might only be 2nd order in these cases. A similar model considered in [36] assumes a one-dimensional phenotype space, so their observed transition can be predicted to be second-order.

control parameter, the transition is conjectured to be continuous (in the limit $\sigma \to 0$, the strains continuously become more uniform and approach the neutral case). Direct checks of these hypotheses (by e.g. directly testing for hysteresis or measuring quantities like critical exponents) will be difficult to observe directly since in the model, demographic noise, mutation and disorder are all strong (and so in finite systems, the transitions are potentially 'smeared'). We summarize these time scale estimates in a conjectured phase diagram in Fig. 4 in the Main text.

We notice that the stability condition for $p$ (from the left of the transition) and the condition for $\sigma$ (from the right of the transition) coincide when expressed in terms of the population sizes, which are easily measured quantities, so we can define the 'eco-evolutionary temperature:'

$$T \equiv paX_{tot}Y_{tot}$$

where $T \gg 1$ is a necessary condition for stability. Both of our separate estimates give a location for the transition that is consistent with simulations (see Fig. 3 in the Main Text).

Importantly, the extinction transition in our model can be thought of as an ergodicity breaking event. Once the predators go extinct, the prey are 'locked in' to their frequencies at the times of extinction, and will remain at those frequencies for potentially long times (until mutation sufficiently scrambles them). However, if more complicated interactions between the surviving species exist (for instance, such that they exhibit chaotic dynamics [28] through internal predation, strain structure, and potentially spatial structure), a weaker form of ergodicity might persist. This would be a potentially interesting topic for future study.

##### D.3 The effective disorder $\sigma_{eff}$ and the Griffiths phase phenomenon

The stochastic transition between predator extinction and stability is analogous to the spin glass (extinct phase) to paramagnetic (stable phase) transition in disordered models of magnetism. A well-known effect in magnetism is that with disorder, there is a continuous region above the spin glass transition point known as a Griffiths phase. In magnetism, a Griffiths phase is one in which a system does not fully fall into a paramagnetic or ferromagnetic phase at a given time. Rather, it is dominated by macroscopic 'rare regions' that relax slowly, that have some average activity. This effect has been demonstrated to be present in classical nonequilibrium and quantum models [8, 22, 21, 33, 34]. Recently, in an ecological context, a similar phenomenon was described as 'species packing' [6], whereby mean field abundances drop well below 1, so that demographic noise induces strong fluctuations.

We can understand the Griffiths phase phenomenon intuitively in the case of our model. For fixed population sizes $X_{tot}$ and $Y_{tot}$ and for fixed number of genotypes $2^L$, when $2^L \gtrsim X_{tot} \sim Y_{tot}$ there are more possible genotypes than

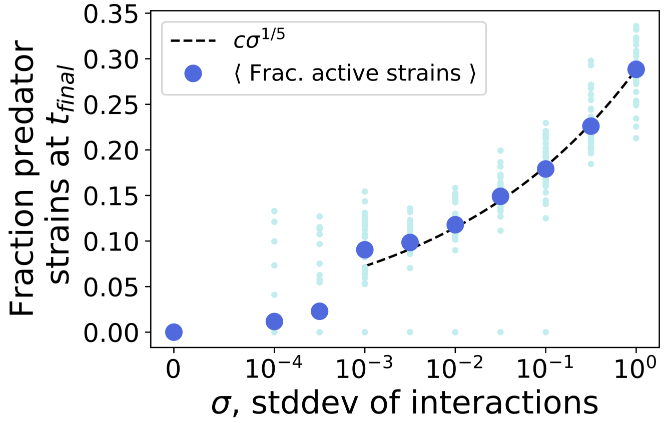


Figure 16: Fraction of active ($Y_k > 0$) predator strains in stochastic simulations (p=0.1) at the final timepoint. As is the case in Griffiths phases in generic nonequilibrium and quantum systems, only a fraction of sites are active even though in the deterministic dynamics all sites are active. The fitted power law exponent is different than that obtained for the total population size.

can exist at once at finite abundance dictated by mean-field, given stochastic effects and the particularities of a single realization of disorder. This means only a subset will exist at finite abundance at a given time, although in the long time limit the average abundances should approach the ones obtained from the deterministic model. In a real predator-prey system, we expect genotype space to be fairly large (much larger than any reasonable population size) so that a potnetially vanishingly small subset of all possible genotypes may exist at a given time.

The presence of a Griffiths phase does not change our arguments for the location of the transition. However, there should be a second transition where the Griffiths phase gives way to the deterministic expectation. But we note that as $\sigma$ gets larger (for fixed $L$), the validity of linear response becomes more suspect since perturbations are no longer necessarily small. Furthermore, once $\sigma$ gets sufficiently large, some rate parameters can become so large to be physically unrealistic models of predator and prey interactions. So for all intents and purposes the Griffiths phase may well extend over a sufficiently large region of the phase diagram that the deterministic dynamics are not representative of any real (i.e. stochastic) system over short times. However, even if this is the case, the deterministic dynamics can still be representative of long time averages.

Finally we comment that the Griffiths phase phenomenon is the likely cause of the discrepancy between the exponent in the stochastic and deterministic dynamics. Generic power laws are common in Griffiths phases due to exponentially large active domains lasting for exponentially distributed times. The convolution of these two factors can lead to a generic exponent [34, 33]. We expect that in the stochastic model, strains are present for times exponentially distributed in their population size so that the response must be averaged over these timescales in addition to being averaged over disorder, leading to a different power than in the deterministic model. However, this effect would need to be verified with a careful analysis of the time-dependent dynamics of the strains, which is unavailable at the current time.

#### D.4 A summary of the stochastic model and some open questions

In the stochastic model, we have used simulations to show that despite having an interaction landscape with underlying standard deviation $\sigma$, the observed standard deviation at a specific time $\sigma_{eff}(t)$ might be much lower than expected. This is due to demographic noise which induces temporary extinctions. However, on sufficiently long timescales, the model realizes an approximately uniform abundance distribution, which allows us to use our analysis of the mean field scenario. Despite a heuristic understanding of the regularizing effect of demographic noise, it would be useful to work out a quantitative understanding of $\sigma_{eff}$ and its dependence on the parameters $a$, $s$, $\sigma$, and $p$. However, such a task will likely require more sophisticated analysis methods than those presented here.

Another important open question can be formulated as 'just how stable is the stable phase?' By approaching the transition from the left, from the extinct phase towards the stable phase, varying $p$, we were able to use relatively simple estimates of the extinction and mutation timescales. However, approaching the transition from the right, from the stable phase towards the extinct phase, what are the relevant scalings of the mutation and extinction timescale? What are the relevant scalings of the extinction and mutation timescale for sufficiently large $\sigma$? Because there are many strains at once, which collectively stabilize each other, might we treat the extinction process with a WKB-like formalism [2]? Will the scalings remain relatively simple functions of the total population size or take some other form that also depends on the number of active strains and their interactions?

Generally, a more rigorous analysis has the potential do provide deeper insights into the phenomenology described here.

### Appendix E Steady population dynamics belie complex strain dynamics

Previous studies have shown that related eco-evolutionary (or epi-evolutionary) models can be mapped onto so-called traveling wave models of population genetics in certain parameter regimes [37, 29, 16]. Using our simulations, we can also access the strain dynamics. So it is natural then to ask whether such a mapping exists for the predator-prey model under consideration.

An essential ingredient of traveling wave models of evolution is that there be a large influx of mutations so that multiple mutations compete with each other or even accumulate within lineages. Qualitatively speaking, in the case of beneficial mutations (n.b. these models have also been analyzed in the context of Muller's ratchet/deleterious mutations), a beneficial mutant grows relative to the rest of the population as low fitness clones are purged and new beneficial mutants are seeded. These processes balance to generate an approximately Gaussian wave profile which travels at fixed speed in fitness space [7, 12].

In order to compare the present model with traveling wave models, we define the prey and predator fitness:

$$g_i^{prey}(t) \equiv b_i - \sum_k (a_{ik} + s_{ik}) Y_k$$

$$g_k^{pred}(t) \equiv -d_i - \sum_i a_{ik} X_i$$

so that the prey fitness is bounded from above:

$$g_i^{prey}(t) \leq b_i$$

and the predator fitness is bounded from below:

$$g_k^{pred}(t) \geq -d_i$$

In the absence of predators, the prey population grows exponentially and in the absence of prey, the predator population decays exponentially. We also neglect the mutational term, which we are assuming to be large enough that there are concurrent mutations but still relatively small (consistent with the case in Fig 17).

We can readily see that the population fitness distribution does not have a limiting shape which stands in stark contrast to the traveling wave picture, cf Fig. 17. In the present model, we see that the most fit genotype will typically have very high fitness, potentially orders of magnitude larger than the mean fitness. This is due to the fact that the fitness of a prey (predator) type depends on the predator (prey) population linearly. If a prey (predator) has an anomalously small (large) interaction with a high abundance predator (prey) type, it will have a relatively large fitness. However, this advantage is only transient as this highly fit type quickly gets pulled into the bulk of the fitness distribution and typically goes extinct as the environment changes. Because of

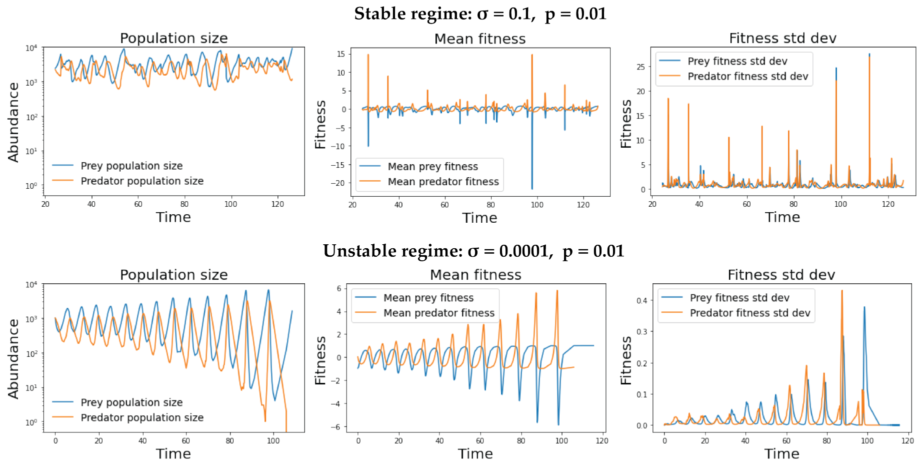


Figure 17: Abundance, mean fitness, and standard deviation of the fitness distribution in the stable regime and in the extinct regime, plotted for stochastic simulations. In the extinct regime, the dynamics of the fitness distribution is smooth, whereas in the stable regime it is intermittent. This is markedly different from traveling wave models of adaptation, where a stable fitness distribution forms.

this 'leapfrog' character, it is difficult to project these high-dimensional dynamics down onto the low-dimensional space of traveling wave models as Yan, et al did for a related model [37].

Instead, we can describe these dynamics in the language of eco-evolutionary feedback. We have established that without sufficient phenotypic variation, either the predator population or both populations will go extinct. However, when there is sufficient variation accessible by mutation, a prey which is anomalously free from predation will be born and will rise exponentially at its growth rate $b_i$. As it rises, its strongest predator will also start to rise at the point where $a_{ik}X_i \sim d_k$. The two focal types will then peak and start to decline, according to the coupled equations:

$$\begin{aligned} \dot{X}_i &\approx X_i(b_i - (a_{ik} + s_{ik})Y_k) \\ \dot{Y}_k &\approx Y_k(-d_k + a_{ik}X_i) \end{aligned}$$

Eventually $X_i$ reaches low enough abundance (and likely goes extinct because there are sufficiently many weakly interacting predators) that $Y_k$ declines at rate $d_k$ until it is also extinct. This whole process happens on a timescale such that as the populations are declining as a whole, a new highly fit prey type is seeded. This results in the observed 'breathing' fitness distribution that spreads across genotype space in potentially complex ways.

In Fig. 17, we see that in the unstable weak selection regime, the mean and standard deviation smoothly oscillate, while in the stable regime the mean and standard deviation show spiky, almost intermittent behavior, with spikes corresponding to the invasion of highly fit types. However these spikes are sufficiently short-lived that they do not distort the total population size by much. A further investigation of the population genetic characteristics of the present model is an interesting avenue for future study.

### Appendix F Relation to other model variants

#### F.1 Correlated landscapes

##### F.1.1 Simulations

We simulated a simple version of the model with phylogenetic correlations by explicitly modeling the binding energies between predator and prey. We considered two cases.

The first case we consider is one in which the interaction energy is additive:

$$E^a_{ik} = \sum_{\alpha=1}^{L} \epsilon_\alpha(i_\alpha, k_\alpha)$$

We draw the $\epsilon_\alpha(i_\alpha, k_\alpha)$ from normal distributions so that the mean and variance of the rates, $a_{ik} = e^{E^a_{ik}}$, are fixed to be $a$ and $\sigma^2$, respectively (on average). The $s_{ik}$ interactions were defined similarly. As $\sigma$ grows, closely related neighbors become more and more uncorrelated (a single step mutation can be large) so that we see the same transition and same qualitative behavior as the uncorrelated model.

We note that in this additive random model, fewer random variables are drawn in defining the interaction landscape ($4L$ vs $2^{2L}$) so that there is a higher variance among realizations. This is a general property of defining finite correlated landscapes. Furthermore, if we look at the strain level dynamics, we see that they are quite different. For a finite sample, strains are no longer "almost exchangeable," and there are clear "winners" which persist at high abundance for long times (i.e. it makes sense to consider fitness in this model). However, this does not change the population level dynamics since enough high fitness types (typically those predator-prey pairs with small interactions and high population size) exist at a time so that there is still a stabilizing effect due to intraspecific competition.

The second case we consider is an energy with 'pairwise' correlations:

$$E^a_{ik} = \sum_{\alpha,\beta=1}^{L} \epsilon_{\alpha,\beta}(i_\alpha, i_\beta, k_\alpha, k_\beta)$$

We draw the $\epsilon_{\alpha,\beta}(i_\alpha, i_\beta, k_\alpha, k_\beta)$ from normal distributions so that the mean and variance of the rates, $a_{ik} = e^{E^a_{ik}}$, are fixed to be (on average) $a$ and $\sigma^2$, respectively. The $s_{ik}$ interactions were defined similarly. When comparing this to the additive model, we see that there are still some "winners," which stay at high abundance for long times. However, the gap between them and the low frequency types has closed a fair amount.

We note that in the additive and pairwise random models, fewer random variables than the uncorrelated genotype model are drawn in defining the interaction landscape ($4L$ vs $16L^2$ vs $2^{2L}$) so that there is a higher sample-to-sample variance among realizations of the random interactions. This is a general property of finite correlated landscapes. Furthermore, if we look at the strain level dynamics, we see that they are quite different. For a finite sample, strains are no longer "almost exchangeable," and there are clear "winners" which persist at high abundance for long times (e.g. it could potentially make sense to define a scalar fitness in such a model). However, this does not change the population level dynamics since enough high fitness types (typically those predator-prey pairs with small interactions and high population size) exist at a time so that there is still a stabilizing effect due to intraspecific competition. And still, in the thermodynamic limit, we expect stability via the same mechanism described above.

We could increase the number of random draws by using more complicated correlation structure. But for our purposes, since we see the same phenomenon-logy (e.g. stabilization) in the additive and pairwise models, we expect it to be present for less-correlated (i.e. higher order) models. These less-correlated models might include, for instance, NK models with $N = L$ and $1 < K < L-1$. Our additive, pairwise and uncorrelated models are the $K = 0$, $K = 1$ and $K = L-1$ cases, respectively. Other models that might be interesting to consider include the "Rought Mt Fuji" model [23] and the Gaussian model of Agarwala and Fisher [1]. However, for the latter, a new simulation scheme would need to be implemented since ours depends on declaring the interaction landscape ahead of time.

We note that there is interesting transient behavior encoded in the model, especially when initialized with a single strain. We see that populations can take a long time to reach the steady state and are much more likely to go extinct by chance when initialized this way. This is similar to what was seen with the fixed $\sigma$, $p$-varying simulation results in the main text. Furthermore, it is interesting that the final (stable) population sizes reached in the correlated models are typically much larger than those reached in the uncorrelated model. This is likely due to the fact that individual types from a finite sample can have much smaller interaction rates in the correlated model than those drawn from an uncorrelated distribution. We leave exploring the details of the additive and other correlated models (especially their asymptotic behavior on infinitely large genotype spaces) as an interesting avenue for future work.

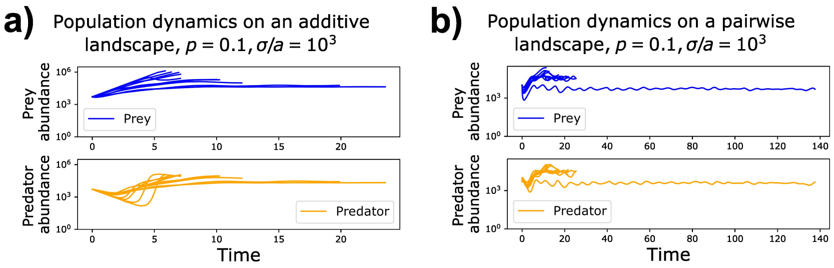


Figure 18: Population dynamics on a.) an additive landscape and b.) a pairwise landscape for $a = 10^{-3}$, $\sigma = 1$, $p = 0.1$. We see similar behavior as the uncorrelated case, where at long times the population seems to go to a fixed point much higher than the fixed point associated with the average interaction strength. However, since it is difficult to control the mean and variance of a finite sample of such a highly correlated landscape, there is a relatively high variance of the populations' final abundances. Populations were initialized with some fraction of strains at finite abundance with total abundance near their deterministic fixed point values. Occasionally, there is extinction due to initial transients. However, this seems to be relatively rare.

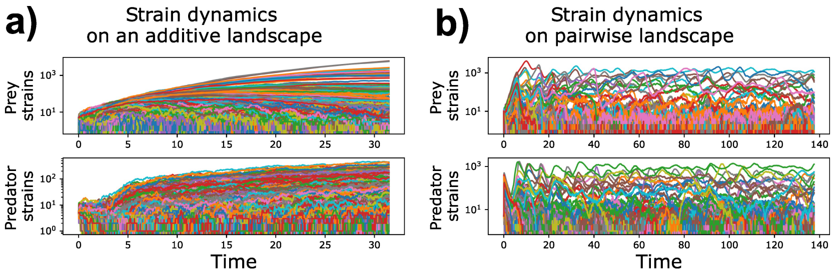


Figure 19: Strain dynamics on a.) an additive landscape and b.) a pairwise landscape for $a = 10^{-3}$, $\sigma = 1$, $p = 0.1$. We see similar behavior as the uncorrelated case, where the strains fluctuate rapidly and over a wide range. However, with correlations, there are clear "winners" among the set of all possible strains.

##### F.1.2 The effect of correlations on the transition

Correlations in interactions space enter into the diffusion length, $l_D$ described in the Section D of this Supplement. In the limit of no correlations (which we primarily discuss in this work), this length is $l_D \sim 1$ in the thermodynamic limit (there is high probability of hitting a beneficial mutation in a single step). This length will increase as the range of the correlations increases. In the case of perfectly correlated interactions, this length is infinite – there are no beneficial mutations to be found no matter how far one mutates. This corresponds to the neutral case in our model, in which there is relatively rapid extinction.

However, in real systems (where we may not be sufficiently close to the thermodynamic limit or where correlations might be long-ranged), this length can be some finite intermediate value – i.e. there are certainly correlations in interaction space, but there are still large jumps that are accessible by few mutations [15, 5]. This can be ameliorated specifically in viral systems, because even if there are correlations in interaction space, this can be compensated by increasing burst size, or by having a sufficiently wide burst size distribution. This will contribute to the effective diffusion constant and bring the diffusion time down. An intuitive way to think about this case is that having a large burst size allows a predator to sample interaction space deeply enough to overcome fitness valleys brought about by correlations.

#### F.2 Relation between the $p$-model and the $\mu$-model

In the main text, we describe a model in which mutations are coupled to births so that they naturally occur on the same timescale as births. However, traditional population genetic analysis has assumed that mutation occurs sufficiently rarely that it occurs *independently of the underlying birth-death process*. The key operational difference is that this results in different classes of diffusion in mutational space.

The constant rate mutational operator is given by:

$$\frac{\mu}{L}\sum_{j\in\mathcal{S}_\rangle}(X_j - X_i)$$

which is the appropriate operator for prey mutations (with $\mu = p$) and for mutations that do not come about as a result of replication machinery. However, since predators are born as a result of interactions, a mutation *probability* operator is more appropriate, though significantly more complicated since it depends on the form of the interactions. For an interaction with a rate given by $F_i(\vec{X})X_i$ which results in a birth (and thus a potential mutation), the appropriate diffusion operator is given by:

$$\frac{p}{L}\sum_{j\in\mathcal{S}_\rangle}(F_j(\vec{X})X_j - F_i(\vec{X})X_i)$$

In a situation where there is sufficient homogeneity or in one where we can use the averaging scheme from DMFT, we can make the identification:

$$\mu = p\langle F_i\rangle_t$$

as we did when estimating the phase boundary. However, there is a caveat that the time average $\langle F_i\rangle_t$ should be carried out in the correct phase.

In cases in which there are high rates of birth-uncorrelated mutations, $\mu \gtrsim 1$ in the parameter regime studied here, we should have similar results to those we have discussed for the $p$ model (i.e. an ergodic stable regime). This sort of scenario might be important, for instance, in natural microbial populations, where horizontal gene transfer can be rampant. In reality however, there is likely a combination of birth uncorrelated and birth correlated mutation. Since birth correlated mutation depends on the interactions, which dominate any other terms in the dynamics, this is likely the more relevant process.

#### F.3 The effect of competition between prey

When left to grow on without predation, bacteria will eventually saturate their environment by consuming all available resources. Therefore it might be important to model this by including a carrying capacity in the prey dynamics. As described in the main text, the results that have been described are valid in the limit where the carrying capacity is sufficiently large when compared to the timescale on which a predation event occurs. For sufficiently large carrying capacity, the populations are 'predation limited.' In this section I will describe the small carrying capacity limit, in which the the prey population is 'resource competition limited' (which indirectly limits the predator population), and I will make a simple argument based on abundance estimates why natural populations are unlikely to exist in this latter limit.

From a stability perspective, the dynamics will not change when there is sufficiently fast evolution. At the mean field level, predator-prey dynamics with a prey carrying capacity possess a stable nontrivial fixed point which we will demonstrate, though this has been shown elsewhere [28, 36, 10]. We can go on to start from the mean field equations for the total population sizes again, except with the addition of the appropriate limitation term:

$$\frac{dX_{tot}}{dt} = \sum_i X_i \left(1 - X_{tot}/K - (a+s)Y_{tot} - \sum_{k=1}^{2^L}(\delta a_{ik} + \delta s_{ik})Y_k\right)$$

$$\frac{dY_{tot}}{dt} = \sum_k Y_k \left(-1 + aX_{tot} + \sum_{i=1}^{2^L}\delta a_{ik} X_i\right)$$

where $K$ is the carrying capacity.

Following the same procedure as in a previous section, applying DMFT and assuming homogeneous strain frequencies (which holds for sufficiently fast evolution, as above, though this is not shown), we obtain the fixed point equations:

$$Y_{tot} = \frac{a - \sigma^2\chi_Y - 1/K}{a(a+s) + \sigma^2\chi_X(\sigma^2\chi_Y + 1/K)}$$

$$X_{tot} = \frac{(a+s) + \sigma^2\chi_X}{a(a+s) + \sigma^2\chi_X(\sigma^2\chi_Y + 1/K)}$$

We see that in the limit $K \to \infty$, we recover our previous results. Notably, there is only a feasible fixed point if the following condition holds:

$$a - \sigma^2\chi_Y > 1/K$$

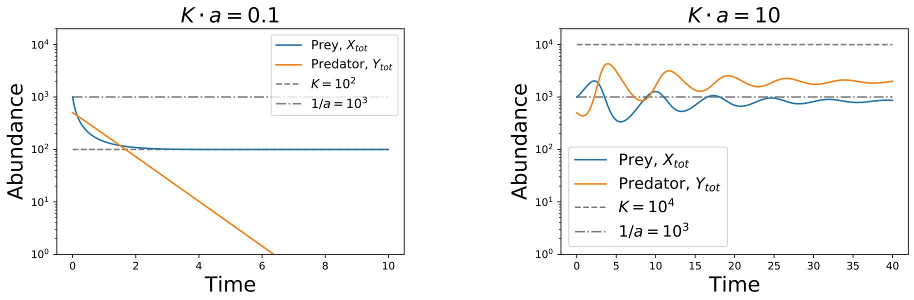


(a) (b)

Figure 20: Mean field dynamics for (a) small and (b) large carrying capacity limits for $\sigma = 0.1$. For sufficiently small carrying capacity, the predator population cannot be sustained. For large carrying capacity, the interactions dominate and set the effective population size of the predator and the prey.

where $\chi_Y$ is again negative when there there is evolution (the carrying capacity does not change the nature of the eco-evolutionary feedback loop). So there is a minimum prey carrying capacity that can support the predator population.

For small enough $\sigma$, we have the approximate inequality constraint for stability:

$$K \cdot a \gtrsim 1$$

which is equivalent to what we described before – the typical timescale for a predation event to occur (given $1/a$) must be less than the carrying capacity (which can be thought of as the typical time for a competition event to occur) so that the ratio $K \cdot a$ is greater than 1 (see Fig. 20). The variance in the rates will simply reduce the right hand side by factor that cannot get larger than $\mathcal{O}(1/a)$ due to the saturation of the response. In the stochastic dynamics, extinctions can still be highly problematic in the purely ecological case even though there is mean-field stability. However, as was demonstrated in [36], in a simplified model, sufficiently fast co-evolution can rescue the populations, with a sharp transition that is analogous to the one described here. Given the form of the mean field equations, a sufficiently small allowable prey carrying capacity, $K \sim 1/a$, can cut off the observed power law growth by regularizing the divergence, but only for a large enough $\sigma_{\text{cutoff}}$. The cutoff scale can be estimated using the scaling form of the response functions and the fixed point equations:

$$\sigma_{\text{cutoff}} \sim p \left( \frac{K}{a+s} \right)^{1/\eta}$$

and can be quite large given $\eta \approx 1/4$. Therefore, the resource-limited regime has a limited domain of relevance, requiring fine-tuning of the carrying capacity to closely match the interaction timescale $1/a$ and potentially very large $\sigma$.

We may ask if such a resource limited regime is a reasonable one to find a natural population. Given the fixed point feasibility constraints and given that natural phage populations are quite large, it is unlikely. In the ocean, phage populations are found to be present at approximately $\sim 10^8$ particles per milliliter of seawater, which is approximately ten times larger than bacteria populations [3, 35]). In the human gut microbiome, estimates are that phage populations reach $\sim 10^{10}$ particles per gram of feces, which is compared to $\sim 10^{11}$ bacterial cells per gram of feces [20, 38]. Given these large numbers and the above constraint on stability, it is expected then that $a - \sigma^2\chi_Y \gg 1/K$. Therefore natural predator-prey populations are more consistent with predator limitation rather than resource limitation under the current model assumptions. However, considering the multitude of other processes that might contribute to such natural population dynamics, it is important to note that this is a very rough estimate and that certain situations might be complicated by other relevant contingencies, such as spatial structure. We discuss a commonly studied type of spatial structure in the next section.

#### F.4 Relation to island models of ecology

Mutation-selection dynamics are analogous to a type of immigration-selection process, though the two processes differ in some key respects. The mutational picture can be mapped onto a high dimensional spatial landscape, where genotypes correspond to islands and mutation acts as migration. However, interactions happen across islands, instead of being localized to a single island. Furthermore, mutation is inherently different from migration in the sense that mutation can be reasonably coupled to the interactions (as we have described in the previous appendix). In situations where mutation is frequent and the population size fluctuates, mutation cannot be simply be decoupled from the population dynamics. When mutation is sufficiently rare (that it can be approximated as regular diffusion), the mapping to spatial models is consistent. In fact, we expect a correspondence between qualitative aspects the stable phase of our model and the stable spatiotemporal chaos phase in [28, 30]. Despite this, there are key differences, specifically that our model is well-mixed and does not require discrete islands, so that there are no refugia for strains to re-invade from. In other words there is no external flux allowing the populations to maintain a finite time-averaged size, but rather this must maintained by consistent beneficial mutants which are internally generated.

We can compare to a simplified spatial structure, by simulating a mainland-island model with constant immigration rate, $\lambda$ (where a population size fixed point can also emerge). The eco-evolutionary model significantly differs from mainland-island models in that the mutation term for a single strain fluctuates wildly between positive values (immigration in) and negative values (emigration out), which average out over long times. In the constant immigration model, there is constant influx of strains maintaining the populations. In addition, in a constant immigration model, extremely large immigration rates are needed to have population stability ($\lambda$ much greater than the interaction term). This ends up changing qualitative aspects of the fixed point, including switching the ordering of the magnitudes of $X_{tot}$ and $Y_{tot}$. In the mutation model, the fixed

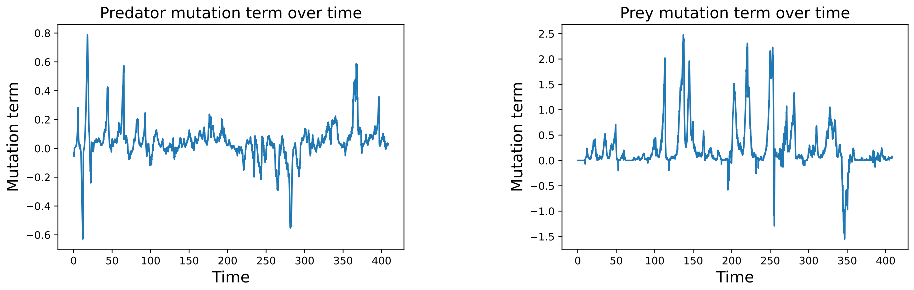


(a) (b)

Figure 21: (a) Typical predator and (b) prey mutation terms over time for $p = 0.1$, $\sigma = 0.01$, for single strains. They fluctuate greatly between positive values (effective immigration in) and negative values (effective emigration out), but generally remain less than or equal to the typical magnitude of the birth rates, death rates and interactions (which in this case is $\approx 1$).

point has $X_{tot} > Y_{tot}$, whereas in a constant immigration model $X_{tot} < Y_{tot}$.

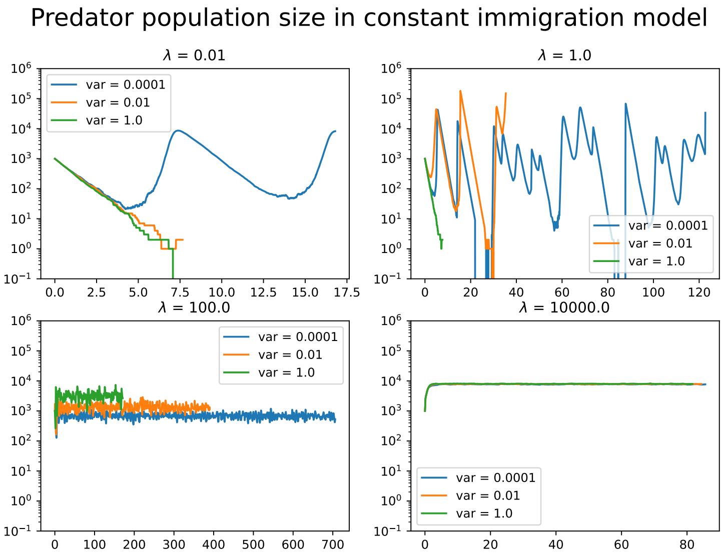


(a)

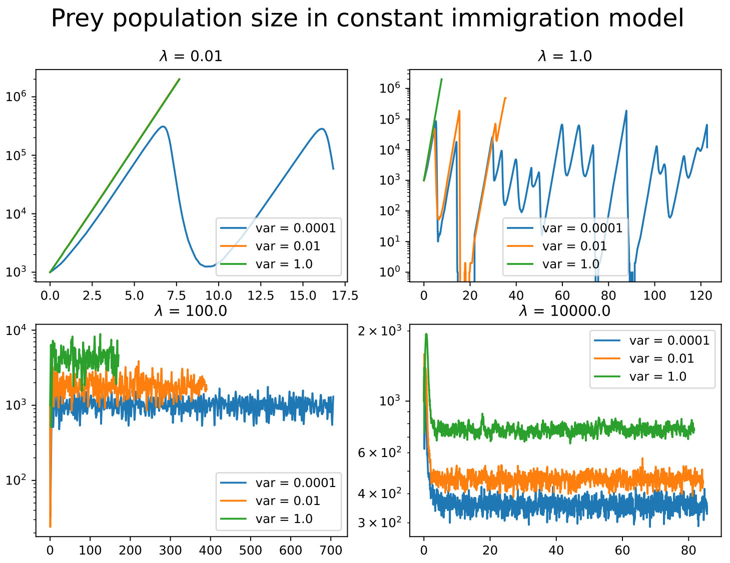


(b)

Figure 22: (a) Predator and (b) prey total population size dynamics for a constant immigration model with varying immigration rates $\lambda$. In the immigration dominated limit ($\lambda \gg 1$), the predator and prey population sizes are inverted from those in the mutation model.

### References


[1] Atish Agarwala and Daniel S Fisher. “Adaptive walks on high-dimensional fitness landscapes and seascapes with distance-dependent statistics”. In: *Theoretical population biology* 130 (2019), pp. 13–49.

[2] Michael Assaf and Baruch Meerson. “WKB theory of large deviations in stochastic populations”. In: *Journal of Physics A: Mathematical and Theoretical* 50.26 (2017), p. 263001.

[3] Øivind Bergh et al. “High abundance of viruses found in aquatic environments”. In: *Nature* 340.6233 (1989), pp. 467–468.

[4] Guy Bunin. “Ecological communities with Lotka-Volterra dynamics”. In: *Physical Review E* 95.4 (2017), p. 042414.

[5] Brian C Cunningham and James A Wells. “High-resolution epitope mapping of hGH-receptor interactions by alanine-scanning mutagenesis”. In: *Science* 244.4908 (1989), pp. 1081–1085.

[6] Jonas Denk and Oskar Hallatschek. “Self-consistent dispersal puts tight constraints on the spatiotemporal organization of species-rich metacommunities”. In: *Proceedings of the National Academy of Sciences* 119.26 (2022), e2200390119.

[7] Michael M Desai and Daniel S Fisher. “Beneficial mutation–selection balance and the effect of linkage on positive selection”. In: *Genetics* 176.3 (2007), pp. 1759–1798.

[8] Daniel S Fisher. “Critical behavior of random transverse-field Ising spin chains”. In: *Physical review b* 51.10 (1995), p. 6411.

[9] Tobias Galla. “Two-population replicator dynamics and number of Nash equilibria in matrix games”. In: *EPL (Europhysics Letters)* 78.2 (2007), p. 20005.

[10] Narendra S Goel, Samaresh C Maitra, and Elliott W Montroll. “On the Volterra and other nonlinear models of interacting populations”. In: *Reviews of modern physics* 43.2 (1971), p. 231.

[11] Nigel Goldenfeld. *Lectures on phase transitions and the renormalization group*. CRC Press, 2018.

[12] Benjamin H Good et al. “Distribution of fixed beneficial mutations and the rate of adaptation in asexual populations”. In: *Proceedings of the National Academy of Sciences* 109.13 (2012), pp. 4950–4955.

[13] Peter Grassberger. “On phase transitions in Schlögl’s second model”. In: *Nonlinear Phenomena in Chemical Dynamics*. Springer, 1981, pp. 262–262.

[14] Hans-Karl Janssen. “On the nonequilibrium phase transition in reaction-diffusion systems with an absorbing stationary state”. In: *Zeitschrift für Physik B Condensed Matter* 42.2 (1981), pp. 151–154.

[15] Thuy-Lan V Lite et al. "Uncovering the basis of protein-protein interaction specificity with a combinatorially complete library". In: *Elife* 9 (2020), e60924.

[16] Jacopo Marchi et al. "Antigenic waves of virus–immune coevolution". In: *Proceedings of the National Academy of Sciences* 118.27 (2021), e2103398118.

[17] Paula Villa Martin et al. "Eluding catastrophic shifts". In: *Proceedings of the National Academy of Sciences* 112.15 (2015), E1828–E1836.

[18] Robert M May. "Will a large complex system be stable?" In: *Nature* 238.5364 (1972), pp. 413–414.

[19] Marc Mézard, Giorgio Parisi, and Miguel Angel Virasoro. *Spin glass theory and beyond: An Introduction to the Replica Method and Its Applications.* Vol. 9. World Scientific Publishing Company, 1987.

[20] Mohammadali Khan Mirzaei and Corinne F Maurice. "Ménage à trois in the human gut: interactions between host, bacteria and phages". In: *Nature Reviews Microbiology* 15.7 (2017), pp. 397–408.

[21] Paolo Moretti and Miguel A Muñoz. "Griffiths phases and the stretching of criticality in brain networks". In: *Nature communications* 4.1 (2013), pp. 1–10.

[22] Miguel A Munoz et al. "Griffiths phases on complex networks". In: *Physical review letters* 105.12 (2010), p. 128701.

[23] Johannes Neidhart, Ivan G Szendro, and Joachim Krug. "Adaptation in tunably rugged fitness landscapes: the rough Mount Fuji model". In: *Genetics* 198.2 (2014), pp. 699–721.

[24] Manfred Opper and Sigurd Diederich. "Phase transition and 1/f noise in a game dynamical model". In: *Physical review letters* 69.10 (1992), p. 1616.

[25] Otso Ovaskainen and Baruch Meerson. "Stochastic models of population extinction". In: *Trends in ecology & evolution* 25.11 (2010), pp. 643–652.

[26] Matthew Parker and Alex Kamenev. "Extinction in the Lotka-Volterra model". In: *Physical Review E* 80.2 (2009), p. 021129.

[27] Matthew Parker and Alex Kamenev. "Mean extinction time in predator-prey model". In: *Journal of Statistical Physics* 141.2 (2010), pp. 201–216.

[28] Michael T Pearce, Atish Agarwala, and Daniel S Fisher. "Stabilization of extensive fine-scale diversity by ecologically driven spatiotemporal chaos". In: *Proceedings of the National Academy of Sciences* (2020).

[29] Igor M Rouzine and Ganna Rozhnova. "Antigenic evolution of viruses in host populations". In: *PLoS pathogens* 14.9 (2018), e1007291.

[30] Felix Roy et al. "Can endogenous fluctuations persist in high-diversity ecosystems?" In: *arXiv preprint arXiv:1908.03348* (2019).

[31] Felix Roy et al. "Complex interactions can create persistent fluctuations in high-diversity ecosystems". In: *PLoS computational biology* 16.5 (2020), e1007827.

[32] Felix Roy et al. "Numerical implementation of dynamical mean field theory for disordered systems: Application to the Lotka–Volterra model of ecosystems". In: *Journal of Physics A: Mathematical and Theoretical* 52.48 (2019), p. 484001.

[33] Federico Vazquez et al. "Temporal griffiths phases". In: *Physical review letters* 106.23 (2011), p. 235702.

[34] Thomas Vojta. "Rare region effects at classical, quantum and nonequilibrium phase transitions". In: *Journal of Physics A: Mathematical and General* 39.22 (2006), R143.

[35] K Eric Wommack and Rita R Colwell. "Virioplankton: viruses in aquatic ecosystems". In: *Microbiology and molecular biology reviews* 64.1 (2000), pp. 69–114.

[36] Chi Xue and Nigel Goldenfeld. "Coevolution maintains diversity in the Stochastic "Kill the Winner" Model". In: *Physical review letters* 119.26 (2017), p. 268101.

[37] Le Yan, Richard A Neher, and Boris I Shraiman. "Phylodynamic theory of persistence, extinction and speciation of rapidly adapting pathogens". In: *Elife* 8 (2019), e44205.

[38] Michele Zuppi et al. "Phages in the Gut Ecosystem". In: *Frontiers in Cellular and Infection Microbiology* (2022), p. 1348.
